# RNA binding proteins Smaug and Cup induce CCR4-NOT-dependent deadenylation of the *nanos* mRNA in a reconstituted system

**DOI:** 10.1101/2022.05.11.491288

**Authors:** Filip Pekovic, Christiane Rammelt, Jana Kubíková, Jutta Metz, Mandy Jeske, Elmar Wahle

**Affiliations:** Institute of Biochemistry and Biotechnology and Charles Tanford Protein Center, Martin Luther University Halle-Wittenberg, Kurt-Mothes-Strasse 3a, 06120 Halle, Germany; Heidelberg University Biochemistry Center (BZH), Im Neuenheimer Feld 328, 69120 Heidelberg, Germany

**Author notes:** Address correspondence to EW; Correspondence may also be addressed to MJ. RNA Biology Laboratory, Center for Cancer Research, National Cancer Institute, 1050 Boyles Street, Frederick, MD 21702, U.S.A.

## Abstract

Posttranscriptional regulation of the maternal *nanos* mRNA is essential for the development of the anterior – posterior axis of the *Drosophila* embryo. The *nanos* RNA is regulated by the protein Smaug. Binding to Smaug recognition elements (SREs) in the *nanos* 3’-UTR, Smaug nucleates the assembly of a larger repressor complex including the eIF4E-T paralog Cup and five additional proteins. The Smaug-dependent complex represses translation of *nanos* and induces its deadenylation by the CCR4-NOT deadenylase. Here we report an *in vitro* reconstitution of the *Drosophila* CCR4-NOT complex and Smaug-dependent deadenylation. We find that Smaug by itself is sufficient to cause deadenylation by the *Drosophila* or human CCR4-NOT complexes in an SRE-dependent manner. CCR4-NOT subunits NOT10 and NOT11 are dispensable, but the NOT module, consisting of NOT2, NOT3 and the C-terminal part of NOT1, is required. Smaug interacts with the C-terminal domain of NOT3. Both catalytic subunits of CCR4-NOT contribute to Smaug-dependent deadenylation. Whereas the CCR4-NOT complex itself acts distributively, Smaug induces a processive behavior. The cytoplasmic poly(A) binding protein (PABPC) has but a minor effect on Smaug-dependent deadenylation. Among the additional constituents of the Smaug-dependent repressor complex, Cup also facilitates CCR4-NOT-dependent deadenylation, both independently and in cooperation with Smaug.

## INTRODUCTION

Poly(A) tails decorate the 3’ ends of almost all eukaryotic mRNAs. Long poly(A) tails are appended to newly made mRNAs in the cell nucleus (Alles et al., 2021; Brawerman, 1981; Eisen et al., 2020; Kühn et al., 2017; Sawicki et al., 1977) and then gradually shortened by 3’ exonucleases in the cytoplasm (Eisen et al., 2020; Goldstrohm and Wickens, 2008; Wahle and Winkler, 2013; Wilson and Treisman, 1988). Poly(A) tail shortening (deadenylation) can serve two regulatory purposes: First, deadenylation initiates the decay of most mRNAs and is a prerequisite for their complete degradation, which occurs mostly by hydrolysis of the 5’ cap followed by 5’-3’ degradation (Charenton and Graille, 2018; Eisen et al., 2020; Muhlrad et al., 1994). The rates of deadenylation are specific for different mRNAs and are the major determinants of mRNA half-life (Decker and Parker, 1993; Eisen et al., 2020; Wilson and Treisman, 1988). Second, deadenylation can contribute to the regulation of translation. Poly(A) tails stimulate the initiation of translation via a protein-mediated interaction with the mRNA 5’ end (Park et al., 2011; Tarun and Sachs, 1996). During oocyte maturation and early embryonic development of animals, it is not just the presence but the length of the poly(A) tail that affects the rate of translation, as initially shown by studies of individual mRNAs in several species (Richter, 2000). Transcriptome-wide analyses confirmed a strong correlation between long tails and high translation efficiency at early embryonic stages, but not in non-embryonic cells (Chang et al., 2014; Eichhorn et al., 2016; Legnini et al., 2019; Lim et al., 2016; Lima et al., 2017; Subtelny et al., 2014; Xiang and Bartel, 2021). In oocyte maturation and early development, regulated extension or shortening of poly(A) tails is used as a means for translational activation or inactivation, respectively (Eckmann et al., 2011; Richter, 2000).

Cytoplasmic deadenylation of mRNAs is catalyzed mainly by the heterooligomeric CCR4-NOT complex (Temme et al., 2014; Tucker et al., 2001; Yi et al., 2018). The complex is organized around the huge central NOT1 subunit (**Fig. 1A**). The N-terminal region of NOT1 binds two subunits, NOT10 and NOT11, which may contribute to substrate RNA recognition (Bawankar et al., 2013; Mauxion et al., 2013; Raisch et al., 2019). The MIF4G domain in the middle of NOT1 provides the docking site for the two catalytic subunits: CAF1 (encoded by *Pop2* in Drosophila) is bound directly, whereas CCR4 (*twin*) binds CAF1 (Chen et al., 2021) and thus associates with NOT1 indirectly. CAF40 (*Rcd1;* CNOT9 in humans), which also has affinity for RNA (Garces et al., 2007; Raisch et al., 2019), associates with the CNOT9 binding domain (CN9BD) of NOT1, which neighbors the MIF4G domain on the C-terminal side (Chen et al., 2014b; Mathys et al., 2014). A NOT2·NOT3 heterodimer (NOT2 encoded by *Regena*) bound to a C-terminal fragment of NOT1 is termed the NOT module (Bhaskar et al., 2013; Boland et al., 2013) (earlier work on the structure of CCR4-NOT reviewed by (Wahle and Winkler, 2013)). The subunits without enzymatic activity not only enhance the activity and substrate specificity of the two exonucleases (Pavanello et al., 2018; Raisch et al., 2019; Stowell et al., 2016), but also provide a large surface to interact with numerous effectors of deadenylation. Each effector binds a specific set of mRNAs and promotes their deadenylation. One class of such deadenylation specificity factors are miRNAs, which recruit the CCR4-NOT complex via associated GW182 proteins (Braun et al., 2011; Chekulaeva et al., 2011; Fabian et al., 2011; Wu et al., 2006). The second class are RNA-binding proteins that, like miRNAs, occupy specific binding sites in 3’ UTRs. Well-characterized examples include tristetraprolin (Brooks and Blackshear, 2013; Bulbrook et al., 2018; Fabian et al., 2013; Webster et al., 2019), Pumilio (Enwerem et al., 2021; Goldstrohm et al., 2018; Webster et al., 2019; Wickens et al., 2002) and Roquin (Leppek et al., 2013; Sgromo et al., 2017), all of which interact directly with the CCR4-NOT complex.

**Figure 1.**
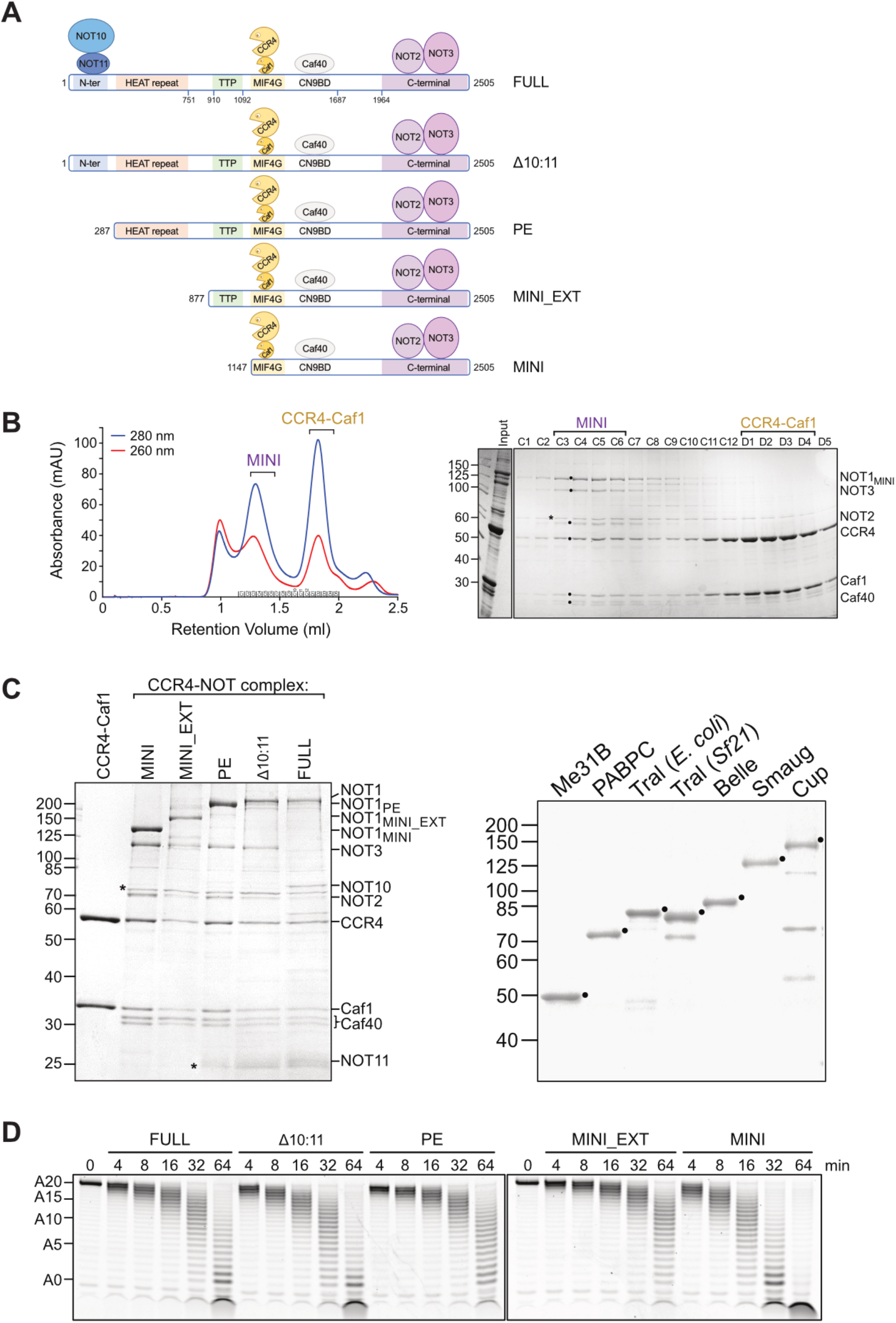
Reconstitution of the *Drosophila melanogaster* CCR4-NOT complex. (A) Scheme of different variants of the *Drosophila* CCR4-NOT complex. Domains and interaction sites of NOT1 are denoted as follows: MIF4G, ‘middle of 4G’ domain; CN9BD, CNOT9 (= CAF40) binding domain; Not11, TTP, Not2/Not3 denote the respective interaction surfaces. NOT1_PC_ and NOT1_PE_ are naturally occurring isoforms. (B) Purification of the ^Dm^CCR4-NOT_MINI_ complex by gel filtration. The complex was first purified by FLAG affinity chromatography and then applied to a Superose 6 column as described in Materials and Methods. Left panel, UV profile of the column; right panel, analysis of relevant fractions by SDS polyacrylamide gel electrophoresis and Coomassie staining. Subunits of the CCR4-NOT complex are labeled with black dots and a contaminating band with an asterisk in the gel. (C) Purified proteins used in this study. Left panel, preparations of different variants of the ^Dm^CCR4-NOT complex. Prominent contaminants are indicated with asterisks. Right panel, purified components of the SRE-dependent repressor complex. Desired polypeptides are marked. Purified proteins were separated on SDS-polyacrylamide gels and stained with Coomassie. (D) Basal activity of different variants of the ^Dm^CCR4-NOT complex. Purified variants of the ^Dm^CCR4-NOT complex (6.25 nM each), were incubated with excess FAM-7mer-A_20_ (50 nM), and aliquots were withdrawn as indicated.

Post-transcriptional regulation, by changes in poly(A) tail length or other means, is essential during early embryonic development of animals: Because the newly formed zygotic genome remains inactive during the first few cell cycles, translation depends on mRNAs that have been synthesized and shelved during oocyte growth and are thus contributed to the developing embryo from the maternal genome via the egg (Colegrove-Otero et al., 2005; Laver et al., 2015; O’Farrell, 2015). For example, regulation of the maternal *nanos (nos*) mRNA is important for the establishment of the anterior-posterior body axis in *Drosophila*. Nanos protein, the determinant for the formation of posterior structures, is made exclusively by translation of a small fraction of *nos* mRNA that is localized at the posterior pole. In contrast, the larger, non-localized fraction of *nos* mRNA is repressed (Gavis and Lehmann, 1992, 1994; Wang and Lehmann, 1991). One repressor of *nos* is the protein Smaug, which binds to two Smaug Response Elements (SREs) in the *nos* 3’ UTR (Dahanukar et al., 1999; Dahanukar and Wharton, 1996; Smibert et al., 1999; Smibert et al., 1996). Smaug also induces *nos* mRNA degradation during the first 2 ½ h of embryo development (Bashirullah et al., 1999; Dahanukar and Wharton, 1996).

Smaug and the related yeast protein Vts1p induce deadenylation of their RNA targets by the CCR4-NOT complex (Aviv et al., 2003; Chen et al., 2014a; Eichhorn et al., 2016; Semotok et al., 2005). Smaug-dependent deadenylation contributes to the large-scale destruction of maternal mRNAs as part of the maternal-to-zygotic transition, which transfers the control over the embryo from the maternal to the zygotic genome (Walser and Lipshitz, 2011). Physical and genetic interactions between Smaug and the CCR4-NOT complex have been observed repeatedly (Götze et al., 2017; Semotok et al., 2005; Temme et al., 2010; Zaessinger et al., 2006), but since the interaction has not been examined with purified proteins, it is unclear whether it is direct. Among the many mRNAs deadenylated under the influence of Smaug is *nos*, which has a short poly(A) tail at steady-state (Eichhorn et al., 2016; Gavis et al., 1996; Jeske et al., 2006; Salles et al., 1994; Zaessinger et al., 2006).

Stimulation of CCR4-NOT-catalyzed deadenylation by specificity factors has so far only been reconstituted with *S. pombe* CCR4-NOT and three different effectors (Stowell et al., 2016; Webster et al., 2019). The data were consistent with a simple ‘tethering’ model in which an individual effector protein associates with a specific RNA sequence and recruits CCR4-NOT by direct interaction. The question how Smaug induces mRNA deadenylation is more complicated, as Smaug does not act on SRE-containing RNAs by itself. Rather, it initiates the assembly of a repressor complex containing six other proteins, all of which are conserved: Cup is an oocyte- and embryo-specific paralog of 4E-T and repressor of translation (Chekulaeva et al., 2006; Götze et al., 2017; Jeske et al., 2011; Nakamura et al., 2004; Nelson et al., 2004; Wilhelm et al., 2003). In tethering experiments in cultured cells, Cup and 4E-T induce deadenylation of the bound mRNA, and the proteins co-precipitate with the CCR4-NOT complex (Igreja and Izaurralde, 2011; Räsch et al., 2020), but a direct interaction has not been demonstrated. A third constituent of the Smaug-dependent repressor complex is Me31B (DDX6 in mammals) (Götze et al., 2017; Jeske et al., 2011; Nakamura et al., 2001). Me31B associates with the MIF4G domain of NOT1 on a surface opposite the CAF1 binding site (Chen et al., 2014b; Mathys et al., 2014; Rouya et al., 2014), but Me31B-dependent deadenylation has, to our knowledge, not been reported. Me31B also binds Cup, suggesting the possibility that Cup-dependent deadenylation might be mediated by Me31B (Kamenska et al., 2016; Nishimura et al., 2015; Ozgur et al., 2015; Waghray et al., 2015). Four additional proteins are part of the Smaug-dependent repressor complex (Götze et al., 2017; Jeske et al., 2011): Trailer hitch (Tral; Lsm14 in mammals), one of several proteins associating with Me31B in a mutually exclusive manner (Brandmann et al., 2018; Tritschler et al., 2009; Tritschler et al., 2008); the translation initiation factor eIF4E, which is bound by Cup (Kinkelin et al., 2012; Nakamura et al., 2004; Nelson et al., 2004; Wilhelm et al., 2003; Zappavigna et al., 2004); the cytoplasmic poly(A) binding protein, PABPC; and the DEAD-box protein Belle (DDX3 in mammals). Among these proteins, PABPC is known to facilitate deadenylation by CCR4-NOT; specifically, PABPC has been reported to promote the activity of CCR4 but inhibit CAF1 (Webster et al., 2018; Yi et al., 2018). However, since PABPC is thought to be bound to poly(A) tails in general, it is unlikely to contribute specifically to SRE-dependent deadenylation. Deadenylation of *nos* is disturbed in *belle* mutants (Götze et al., 2017), but a direct involvement of Belle in deadenylation has not been examined. Tral and eIF4E have not been tested for a role in deadenylation.

In this paper we have used biochemical reconstitution assays to address the question whether Smaug can accelerate deadenylation as an individual protein, by direct recruitment of the CCR4-NOT complex, and/or whether additional components of the Smaug-dependent translation repressor complex are also employed for the purpose of deadenylation. We find that both Smaug and Cup, individually and cooperatively, directly promote deadenylation by the CCR4-NOT complex.

## RESULTS

### Reconstitution of the CCR4-NOT complex from *Drosophila melanogaster*

Reconstitution of five different versions of the *Drosophila* CCR4-NOT complex (**Fig. 1A**) was guided by earlier work on the human complex (Raisch et al., 2019): *^Dm^*CCR4-NOT_FULL_ was composed of full-length versions of all eight subunits (CAF1, CCR4, NOT1-3, CAF40, NOT10 and NOT11). ^*Dm*^CCR4-NOT_Δ10:11_ lacked NOT10 and 11. ^*Dm*^CCR4-NOT_PE_ was similar, but contained the naturally occurring NOT1 splice variant PE, which lacks the 5’ portion of the open reading frame (https://flybase.org). In *^Dm^*CCR4-NOT_MINI_EXT_ NOT1 was further shortened, starting with amino acid 877. *^Dm^*CCR4-NOT_MINI_ contained the shortest NOT1 version, starting at amino acid 1147 and thus missing the tristetraprolin binding site mapped in the mammalian ortholog (Fabian et al., 2013). For the production of these complexes, three MultiBac clones were generated, containing NOT1 and NOT2; NOT3 and CAF40-FLAG; or FLAG-CCR4 and CAF1, respectively (Berger et al., 2004; Fitzgerald et al., 2006). Complexes composed of all six subunits were produced by co-infection of insect cells with the three viruses and purified by Flag affinity-purification followed by gel filtration. For the production of ^*Dm*^CCR4-NOT_FULL_, His_6_-MBP-NOT10 and His_6_-NOT11 were co-expressed in *E. coli*. After co-purification, they were mixed with the Flag-purified six subunit complex, and the assembly was purified by gel filtration. The resulting preparations were pure and generally contained approximately stoichiometric amounts of the subunits (**Fig. 1B; 1C, left panel**). Excess CCR4-CAF1 heterodimer was obtained from the same gel filtration columns (**Fig. 1B; 1C, left panel**). The basal deadenylation activities of these complexes were measured in reactions with a small synthetic RNA substrate (‘FAM 7mer-A_20_’: seven nt ‘body’ plus 20 3’-terminal A residues) (Raisch et al., 2019), carrying a fluorescent label at its 5’ end (**Fig. 1D**). Activities of the larger complexes were similar except that the single CCR4-NOT_MINI_ preparation examined was slightly more active than the other complexes. We conclude that NOT10, NOT 11, and the N-terminal half of NOT1 do not increase the basal activity of CCR4-NOT. In contrast, activity of the CCR4-CAF1 heterodimer was about 2% of that of the MINI complex (**Suppl Fig. 1A**), in qualitative agreement with reports for the *S. pombe* and human CCR4-NOT complexes (Pavanello et al., 2018; Raisch et al., 2019; Stowell et al., 2016). The activity of CCR4-NOT_MINI_ was highest at ~ 1 mM Mg^2+^ and ~ 50 mM potassium acetate (**Suppl. Fig. 1B**).

### Smaug is sufficient to induce SRE-dependent deadenylation by the CCR4-NOT complex

In order to examine Smaug-dependent deadenylation in a fully reconstituted system, we overproduced and purified Smaug and its associates: Smaug, Cup, Me31B and Belle were produced in insect cells by means of baculovirus vectors, PABPC was made in *E. coli,* and Tral was made in either system (**Fig. 1C, right panel**). Cup was the least pure among all proteins. Attempts at further purification were not successful as the protein, after elution from the initial Flag affinity column, did not elute in a defined peak from any other column tested.

Short substrate RNAs were used to assay for Smaug-dependent deadenylation: Most experiments employed the ‘SRE-only’ RNA, which contained two synthetic SREs, either wildtype (SRE^WT^) or with a single inactivating point mutation in each (SRE^MUT^) (Jeske et al., 2006). Both RNAs carried a plasmid-encoded poly(A) tail of some 70 nucleotides. Linearisation of the plasmid DNA for run-off transcription was such that no non-A nucleotides were encoded at the end of the poly(A) tail (see Materials and Methods).

SRE-only RNAs were mixed with Smaug and preincubated to allow complex formation. Addition of ^Dm^CCR4-NOT_MINI_ resulted in rapid deadenylation of the SRE^WT^ RNA during an incubation at 25°C; in the SRE^MUT^ control, the fully deadenylated product appeared only after a lag phase of ~ 16 min and then accumulated at a ~ 50fold lower rate compared to SRE^WT^ (**Fig. 2A**). The ^Dm^CCR4-NOT_FULL_ complex behaved similar to MINI (**Suppl. Fig. 1C**). Three observations indicate that the RNA shortening visible in the gel was due to deadenylation: (1) The reaction product had the size expected for deadenylation. (2) Results similar to those seen with internally labeled RNA in **Fig. 2A** were also obtained with 5’-end-labeled RNA (see below, **Fig. 6B**). (3) The reaction was affected by point mutations in the active sites of the deadenylases CCR4 and CAF1 (see below, **Fig. 4**). As negative controls, Smaug by itself did not catalyze deadenylation, and the CCR4-NOT complex by itself had a barely detectable basal deadenylation activity (**Fig. 2A**). As further negative controls, several other RNA binding proteins, including members of the SRE-dependent repressor complex (see below) and human PTB, a regulator of splicing (Kafasla et al., 2012), did not accelerate deadenylation (**data not shown**). Smaug-dependent deadenylation was carried out in the absence of ATP and was not affected by its addition **(data not shown)**. The reaction was facilitated by the presence of polyethylene glycol (PEG 20,000), presumably due to a macromolecular crowding effect (**Suppl. Fig. S2A**). PEG was therefore routinely included in the deadenylation reaction buffer. The activity of the CCR4-NOT complex itself was not affected by PEG (**data not shown**). miRISC-dependent deadenylation has been reported to be associated with liquid-liquid phase separation (LLPS) (Sheu-Gruttadauria and MacRae, 2018), and LLPS is known to be promoted by crowding reagents like PEG (Alberti et al., 2019). However, by the criterium of centrifugation (Sheu-Gruttadauria and MacRae, 2018), no LLPS was detectable in our deadenylation assays.

**Figure 2.**
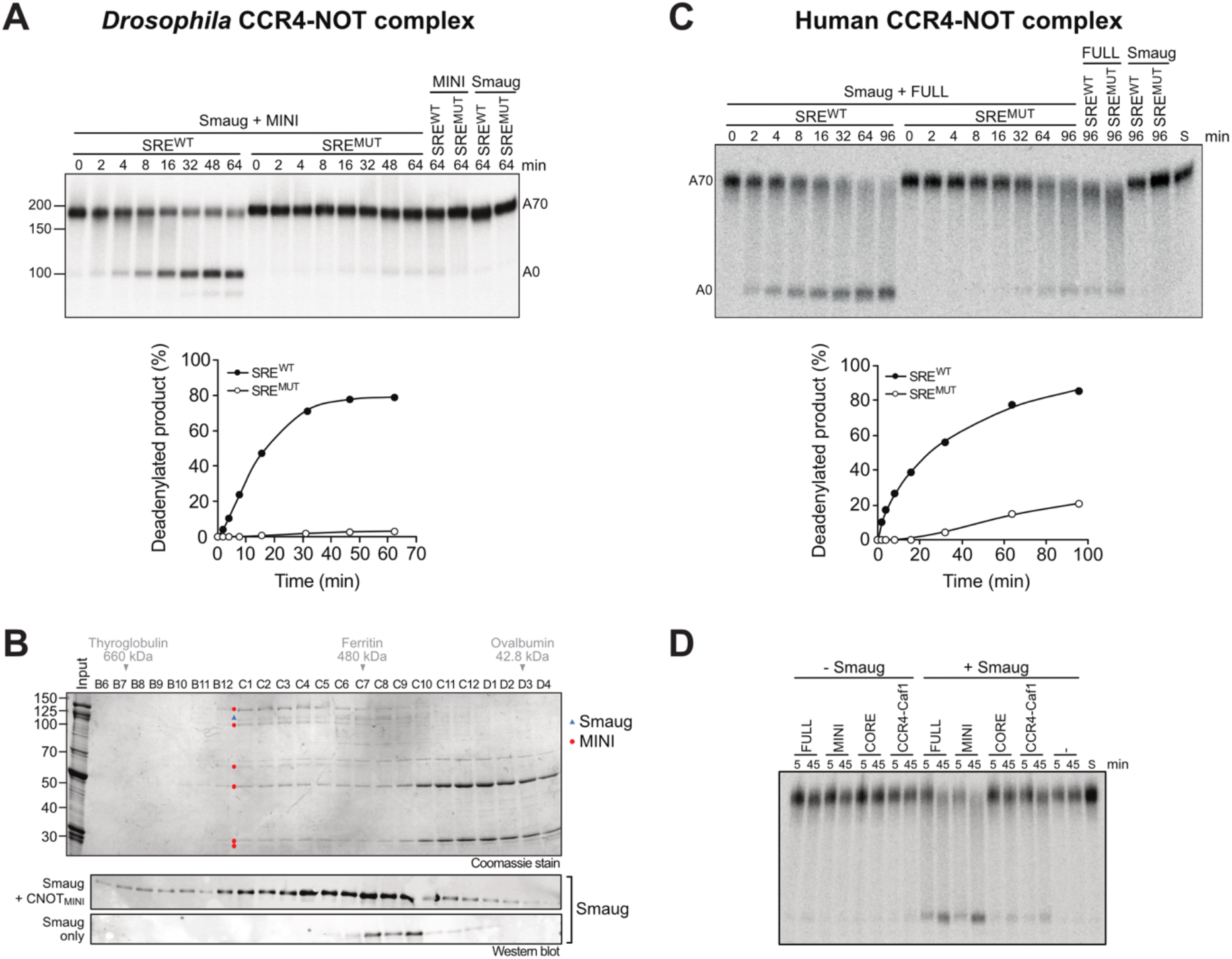
Smaug is sufficient to induce SRE-dependent deadenylation by the CCR4-NOT complex. (A) Smaug induces deadenylation by the ^Dm^CCR4-NOT_MINI_ complex. Radioactively labelled SRE^WT^-A_70_ or SRE^MUT^-A_70_ RNAs (20 nM) were pre-incubated with 80 nM Smaug or deadenylation buffer. Reactions were started by the addition of 2 nM ^Dm^CCR4-NOT_MINI_, and samples were taken and analyzed at the times indicated. Reactions containing only CCR4-NOT or only Smaug were included as controls. The graph at the bottom represents the time-dependent accumulation of fully deadenylated RNA. (B) Gel filtration reveals an association of Smaug with the ^Dm^CCR4-NOT_MINI_ complex. Smaug by itself or mixed with ^Dm^CCR4-NOT_MINI_ was analyzed by gel filtration as described in Material and Methods. Column fractions from the samples containing both CCR4-NOT and Smaug were analyzed by SDS polyacrylamide gel electrophoresis and Coomassie staining (top panel) and by western blotting with an antibody against Smaug (middle panel). Column fractions derived from the Smaug-only sample were only analyzed by western blotting (bottom panel). Elution positions of size markers are indicated above the respective fractions. In the Coomassie-stained gel, Smaug is labeled with a blue triangle, and subunits of the CCR4-NOT complex are labeled with red dots. (C) Smaug induces SRE-dependent deadenylation by the human CCR4-NOT complex. Reactions were carried out as in **Fig. 2A** except that RNA was used at 5 nM, Smaug at 30 nM, and the human CCR4-NOT_FULL_ complex at 10 nM. The graph at the bottom represents the time-dependent accumulation of fully deadenylated RNA. The requirement for a higher concentration of CCR4-NOT compared to the experiment in **Fig. 2A** is at least partially explained by the reaction temperature of 25°C, which is suboptimal for the human complex. These reactions were also carried out in the absence of BSA. (D) The CCR4-NOT ‘MINI’ complex is necessary and sufficient for Smaug-dependent deadenylation. 5 nM ^32^P-SRE^WT^-A_70_ RNA was preincubated with 30 nM Smaug or buffer, then deadenylation was initiated by the addition of the different human CCR4-NOT complexes (10 nM of Full, Mini and Core; 50 nM of CCR4-Caf1). Samples were taken and analyzed at the times indicated.

### Smaug interacts with the NOT module

The data so far demonstrate that Smaug induces SRE- and CCR4-NOT-dependent deadenylation independently of its associated repressor proteins. This would be most easily explained by SRE-bound Smaug recruiting the deadenylase through a direct interaction. Such an interaction was in fact demonstrated by analytical gel filtration (**Fig. 2B**): Smaug alone was recovered in low yields; only small amounts were detectable by western blotting. These eluted in fractions C8 – C10, somewhat ahead of the position expected based on the molecular weight of monomeric Smaug. When Smaug was gel-filtered together with ^Dm^CCR4-NOT_MINI_, recovery was improved, Smaug was detectable by Coomassie staining, and a large fraction of the protein co-eluted with CCR4-NOT in fractions C1 – C6.

Smaug also induced SRE-dependent deadenylation by the human CCR4-NOT complex (**Fig. 2C**). In this experiment, eight subunit ^Hs^CCR4-NOT_FULL_ was used, which is essentially complete except for an N-terminal deletion of NOT11 (Raisch et al., 2019). Several subassemblies of the human CCR4-NOT complex have also been purified (Raisch et al., 2019). ^Hs^CCR4-NOT_MINI_, corresponding to ^Dm^CCR4-NOT_MINI_ except for N-terminal deletions of NOT2 and NOT3, catalyzed Smaug- and SRE-dependent deadenylation with an efficiency comparable to the FULL complex (**Fig. 2D**). CCR4-NOT_CORE_ is a complex further simplified by omission of the NOT module (NOT2, NOT3, and a C-terminal part of NOT1) and thus consists only of CAF1, CCR4, and CAF40 bound to a central NOT1 fragment. CCR4-NOT_CORE_ responded to Smaug very weakly. A CCR4-CAF1 heterodimer behaved similarly (**Fig. 2D**). We conclude that the Smaug interaction surface of the CCR4-NOT complex is conserved between *D. melanogaster* and *H. sapiens* and is at least partially contained in the NOT module.

Interactions between Smaug and individual CCR4-NOT subunits present in the MINI complex were examined by a recently described *in vivo* interaction assay based on intracellular protein relocalization (ReLo assay) (Salgania et al., 2022). For this assay, CCR4-NOT subunits were fused with mCherry and with the pleckstrin homology (PH) domain of rat phospholipase Cδ1 and expressed as bait in *Drosophila* S2R+ cells; the PH domain caused their localization on the plasma membrane. Smaug, as potential prey protein, was labeled with EGFP. Upon coexpression with the bait proteins, an interaction was expected to become visible as the colocalization of the two fluorescence markers on the plasma membrane. With five out of the six bait proteins tested, Smaug remained widely distributed in the cytoplasm. Membrane-bound NOT3, however, caused a clear re-localization of Smaug to the plasma membrane, indicating an interaction (**Fig. 3B**). The Smaug interaction surface of NOT3 was mapped by additional ReLo experiments with swapped tags: NOT3 fragments (**Fig. 3A**), fused to the PH domain and mEGFP, were combined with mCherry-tagged Smaug. The results indicated that Smaug binding is mediated by the C-terminal region of NOT3, which contains the NOT1-interacting region and the NOT box (**Fig. 3A, C**). A split-ubiquitin yeast two-hybrid assay (Stagljar et al., 1998), in which Smaug was used as the bait, confirmed the interaction with NOT3 (**Fig. 3D, left panel**). In this experiment, a NOT1 fragment comprising amino acids 752 – 910 also showed an interaction with Smaug (**Fig. 3D, right panel**). However, this region of NOT1 is not present in the CCR4-NOT_MINI_ complex, which is sufficient for Smaug-dependent deadenylation, whereas it is present in the ^Hs^CCR4-NOT_CORE_ complex, which does not respond to Smaug. Therefore, this region of NOT1 is neither required nor sufficient for a functional interaction with Smaug, and we did not pursue the interaction. The yeast two-hybrid assay confirmed the ability of the C-terminal region of NOT3 to mediate the interaction with Smaug (**Fig. 3E**). The Smaug-NOT3 interaction that we describe here is consistent with the requirement for the NOT module in deadenylation assays (**Fig. 2D**). Mapping of the interaction surface to the C-terminal region of NOT3 is also consistent with the activity of ^Hs^CCR4-NOT_MINI_ in Smaug-dependent deadenylation: In this complex, NOT3 is truncated just upstream of the NOT1-interacting region (Raisch et al., 2019). Attempts to map the complementary interaction surface to a specific domain of Smaug were not successful.

**Figure 3.**
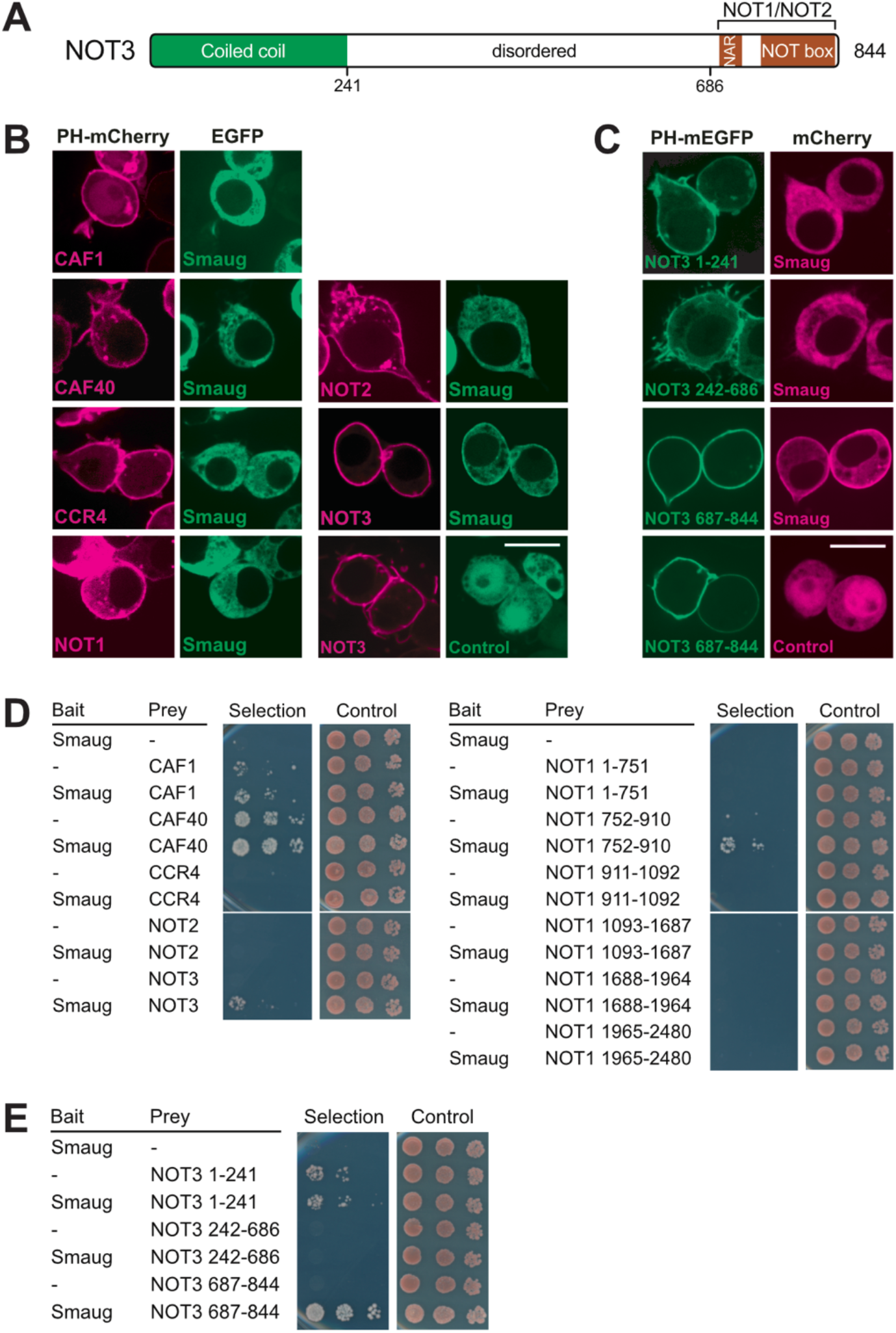
Smaug associates with CCR4-NOT via NOT3. (A) Domain structure of NOT3. NAR, NOT1 anchor region. The NOT box mediates the interaction with NOT2. Borders of fragments used in the interaction assays are indicated at the bottom. (B) Smaug interacts with NOT3 in a relocalization assay. *Drosophila* proteins were fused with PH-mEGFP or mCherry as indicated and transiently coexpressed in *Drosophila* S2R+ cells. After two days, subcellular protein localization was examined by confocal live fluorescence microscopy. Smaug relocalization was detected only with the NOT3 subunit. ‘Control’ indicates a plasmid expressing EGFP only. Scale bar is 10 μm. (C) The Smaug interacting surface is in the C-terminal domain of NOT3 as determined with the ReLo assay. NOT3 fragments were used as bait fusions as indicated. Fluorescent tags were swapped in comparison to **(B).** ‘Control’ indicates a plasmid expressing mCherry only. Scale bar is 10 μm. (D) Smaug interacts with NOT3 in a yeast two-hybrid assay. Split-ubiquitin yeast two-hybrid assays were performed with bait and prey constructs containing *Drosophila* proteins as indicated or no insertion (-). Three 10-fold dilutions of the cells were spotted and imaged after three days of incubation. Selection medium lacked histidine. (E) Yeast two hybrid assay confirms interaction of Smaug with the C-terminal domain of NOT3. The same fragments as in (C) were used as prey fusions in the two-hybrid assay.

### Both CCR4 and Caf1 contribute to Smaug-dependent deadenylation

Earlier experiments employing RNAi or overexpression of catalytically inactive mutants suggested that Caf1 makes the main contribution to the catalytic activity of the CCR4-NOT complex (Arvola et al., 2020; Temme et al., 2010). We have now used the reconstituted complex to examine the relative importance of Caf1 and CCR4. ^Dm^CCR4-NOT_MINI_ was prepared with point mutations in the active sites of either nuclease or the combination of both. In assays employing the FAM 7mer-A_20_ RNA as a substrate, the inactivation of either nuclease subunit had an unexpectedly large effect, reducing the activity of the complex by about 80% (**Fig. 4A**). This behavior was seen with several independent preparations of the enzyme complexes and is most likely explained by either inactive subunit exerting a dominant-negative effect on the active subunit, perhaps by transiently blocking access to the 3’ end. An even stronger effect of the same type has been reported for human CCR4-NOT (Maryati et al., 2015). The combination of mutations in both CCR4 and CAF1 abolished the catalytic activity of the complex (**Fig. 4A**). The Smaug-dependent reaction was also impaired by point mutations in either catalytic subunit. Again, either single mutant had an effect larger than expected, but mutation of CCR4 was slightly more inhibitory than inactivation of CAF1. Combined mutation of both subunits prevented Smaug-dependent deadenylation (**Fig. 4B**). We conclude that, under our experimental conditions, both catalytic subunits of CCR4-NOT make approximately equal contributions to both basal deadenylation and the Smaug-dependent reaction.

**Figure 4.**
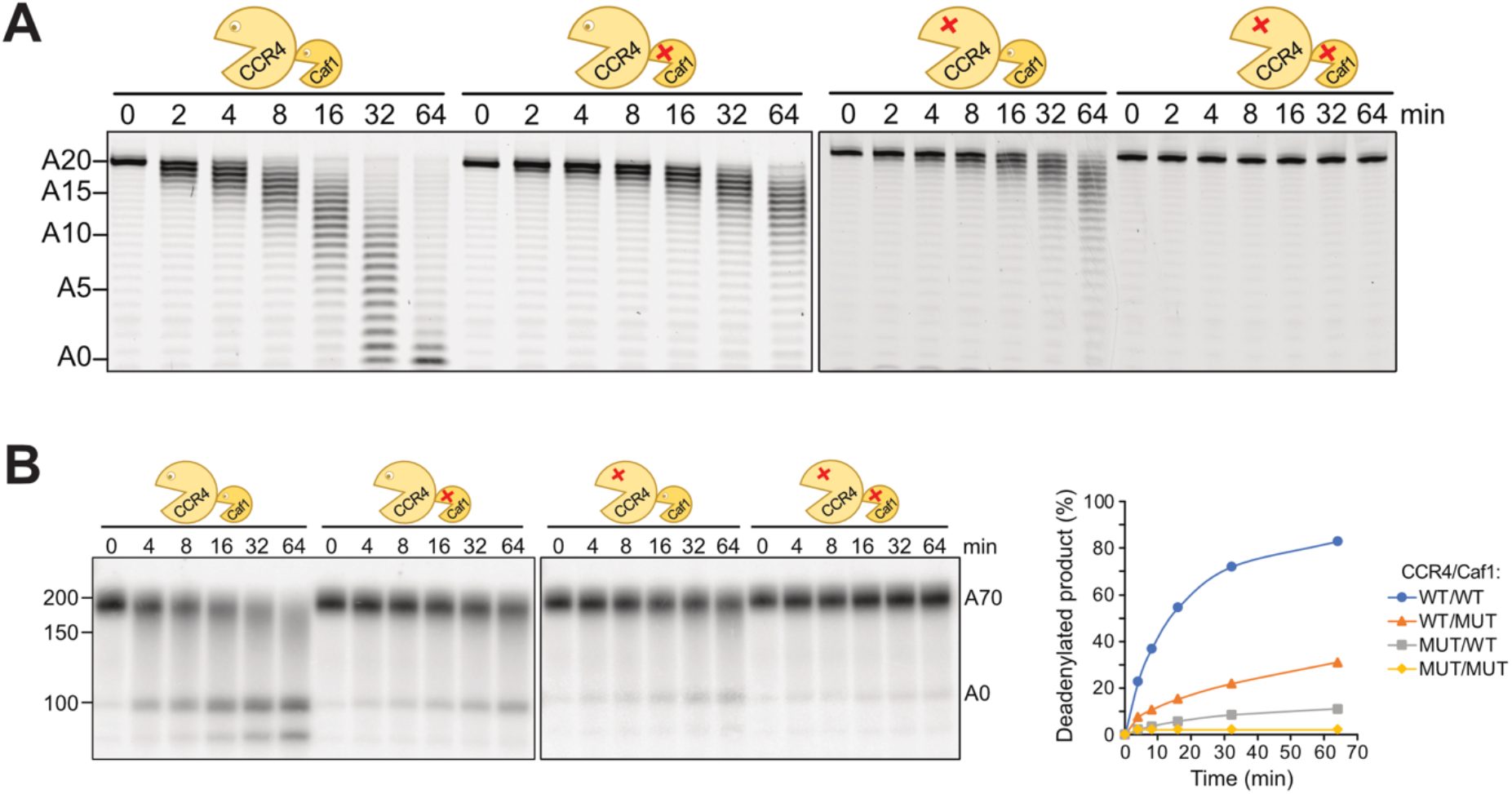
CCR4 and CAF1 make similar contributions to the activity of the ^Dm^CCR4-NOT_MINI_ complex. (A) Inactivation of either catalytic subunit has a similar effect on the basal activity of ^Dm^CCR4-NOT_MINI_. Point mutations in the active sites of CCR4 and Caf1 are depicted in the cartoons. 50 nM FAM 7mer-A_20_ RNA was incubated with the respective enzyme complexes (5 nM) for the times indicated. (B) Inactivation of either catalytic subunit has a similar effect on Smaug-dependent deadenylation. 40 nM Smaug was pre-incubated with 10 nM SRE^WT^-A_70_ substrate RNA, deadenylation was started by the addition of 2 nM ^Dm^CCR4-NOT_MINI_ complex and stopped at the times indicated. The right panel shows a quantification of the fully deadenylated product.

*In vivo,* the substrate for deadenylation is not naked poly(A), but a complex of poly(A) and the cytoplasmic poly(A) binding protein, PABPC. Moreover, PABPC is part of the SRE-dependent repressor complex (Götze et al., 2017). The *S. pombe* and human orthologues stimulate the activities of the cognate CCR4-NOT complexes (Webster et al., 2018; Yi et al., 2018). Thus, we sought to examine the effect of PABPC on deadenylation. With the FAM 7mer-A_20_ RNA, relatively high concentrations of *Drosophila* PABPC were necessary for complete binding (**Suppl. Fig. 3A**), perhaps due to the presence of excess tRNA or partial inactivity of the protein preparation. As expected, the protein stimulated deadenylation by ^Dm^CCR4-NOT_MINI_ both at saturating and sub-saturating concentrations (**Suppl. Fig. 3B**). In a more detailed deadenylation time course, PABPC was initially inhibitory, and the stimulatory effect became apparent only at later times. Size distributions of the RNA were also more heterogeneous in the presence of PABPC (**Fig. 5A**). Presumably, PABPC initially sequesters the 3’ end of the RNA from attack by the nuclease; once the enzyme has gained a foothold on the substrate, PABPC facilitates further deadenylation. Versions of CCR4-NOT carrying inactivating points mutations in either CCR4 or CAF1 responded similarly to PABPC: Both CCR4 and CAF1 were able to degrade poly(A) covered by PABPC, although CCR4 appeared to be slightly more efficient (**Fig. 5B**). With the SREonly RNA carrying a long poly(A) tail, the stimulatory effect of PABPC, in the absence of Smaug, was weak at concentrations that were near saturation for RNA binding (**Fig. 5C**). Note that, under the same conditions, the stimulation by Smaug was much stronger (**Fig. 5D**). The Smaug-dependent degradation of an A70 tail by the CCR4-NOT complex was also moderately stimulated by PABPC (**Fig. 5D**).

**Figure 5.**
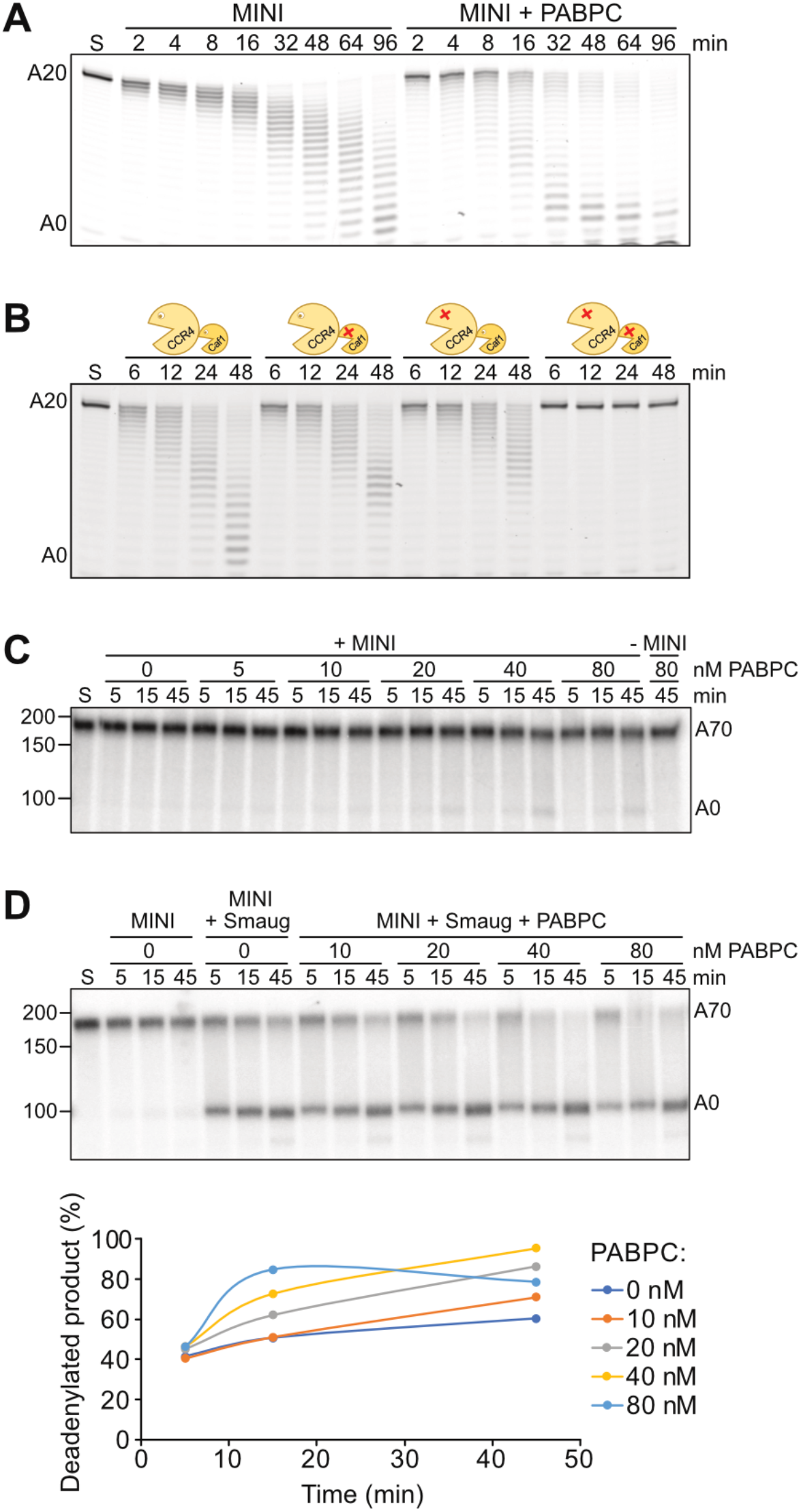
^Dm^CCR4-NOT-dependent deadenylation is stimulated by PABPC. (A) PABPC stimulates the deadenylation of FAM 7mer-A_20_. Substrate RNA (25 nM) was incubated with 200 nM PABPC, and deadenylation was started by the addition of 2.5 nM ^Dm^CCR4-NOT_MINI_. (B) Both CCR4 and CAF1 can degrade a poly(A) tail bound by PABPC. Deadenylation time courses were carried out with wild-type ^Dm^CCR4-NOT_MINI_ or mutant variants as indicated. Reaction conditions were as in (A). (C) PABPC modestly stimulates deadenylation of SREonly-A_70_. SRE^WT^only-A_70_ RNA (5 nM) was deadenylated in the presence of the indicated concentrations of PABPC. ^Dm^CCR4-NOT_MINI_ was used at 0.5 nM. A negative control (last lane) contained 80 nM PABPC in the absence of CCR4-NOT. (D) PABPC modestly stimulates SRE-dependent deadenylation. SRE^WT^only-A_70_ RNA (5 nM) was first pre-incubated with Smaug (30 nM) or buffer for 20 min, then the indicated amounts of PABPC or buffer were added, and the incubation was continued for another 20 min. Finally, deadenylation was started by the addition of ^Dm^CCR4-NOT_MINI_ (0.5 nM).

### Smaug makes deadenylation processive

Accessory factors often boost the activity of nucleic acid-polymerizing or -degrading enzymes by increasing their processivity (Bienroth et al., 1993; Stukenberg et al., 1991). A deadenylase can be judged to be processive by two criteria: First, under conditions of substrate excess, largely or completely deadenylated products will co-exist with untouched substrate because a processive enzyme will act repeatedly, without dissociation, on the small fraction of substrate to which it is first bound, leaving the rest for later rounds. A completely distributive enzyme, in contrast, will remove single nucleotides from random RNA molecules and thus shorten an excess of substrates in a synchronous manner through multiple rounds of association and dissociation. Second, if a poly(A) tail is completely degraded without intermittent dissociation of the enzyme, the rate at which an individual poly(A) tail is shortened will be independent of the concentration of the processive nuclease or its ratio to substrate. Thus, the time at which the deadenylation end product is first seen will be independent of the nuclease concentration; only the amount of end product present at this time will increase with increasing nuclease concentration. In contrast, for a distributive enzyme the time required for the deadenylation end product to appear will be shorter with higher enzyme concentrations as a high concentration will drive the association of enzyme with substrate preceding every catalytic event.

The processivity of the CCR4-NOT complex by itself was first assessed in reactions with the FAM 7mer-A_20_ RNA. As shown in **Fig. 6A**, fully deadenylated products did not co-exist with untouched substrate and became visible only at later time points when all of the substrate had already been shortened to a significant extent. Also, fully deadenylated product first became visible at earlier time points in proportion with increasing enzyme concentration; in other words, the rate of shortening of an individual poly(A) tail was dependent on the nuclease concentration. By both criteria, the activity of CCR4-NOT with this substrate was distributive.

**Figure 6.**
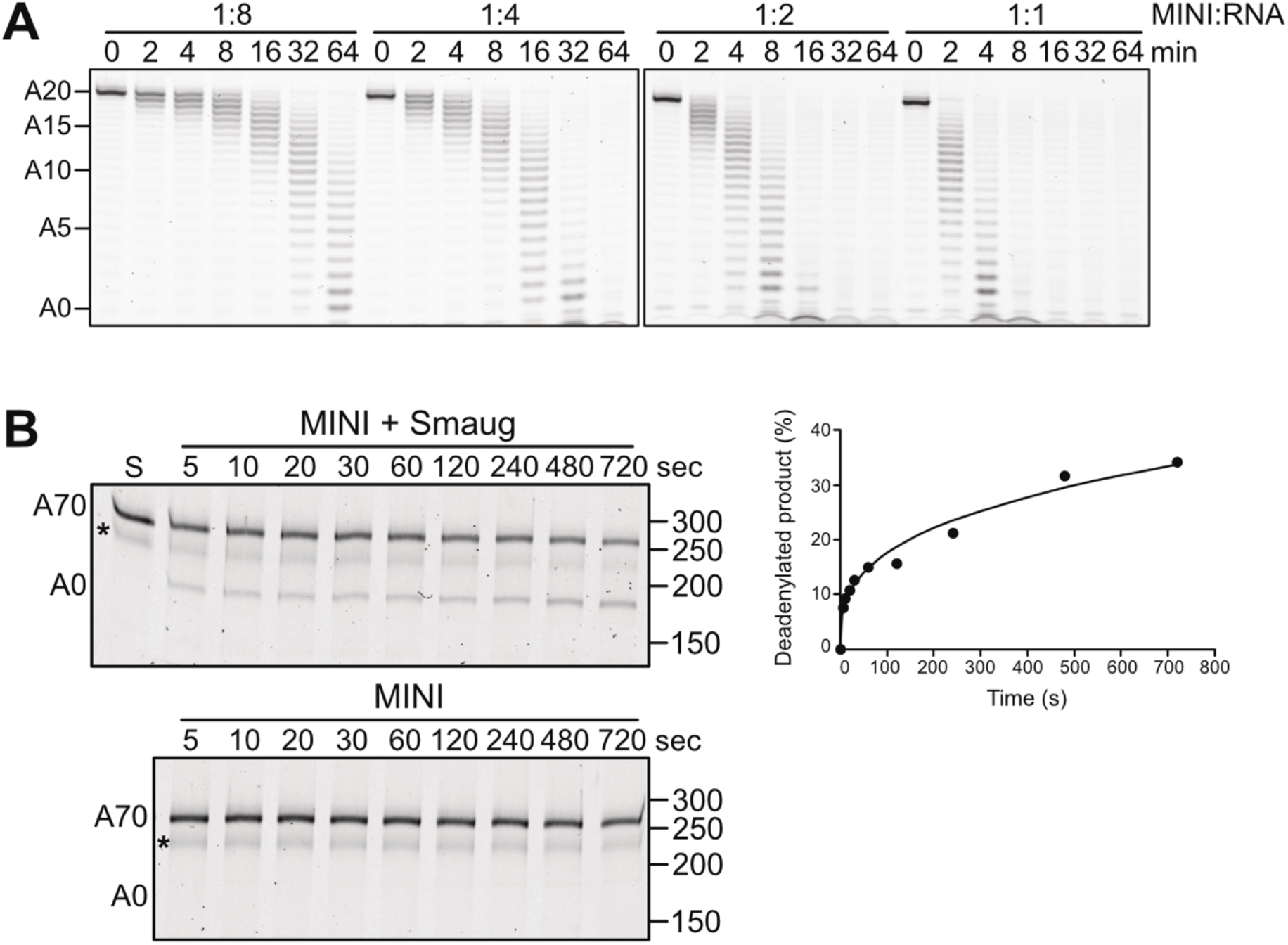
Smaug makes the CCR4-NOT complex processive. (A) The ^Dm^CCR4-NOT_MINI_ complex is distributive on its own. A constant concentration of the FAM 7mer-A_20_ RNA (50 nM) was incubated with 6.25, 12.5, 25 or 50 nM ^Dm^CCR4-NOT_MINI_, resulting in the molar ratios indicated. Aliquots were withdrawn at the time points indicated. (B) The ^Dm^CCR4-NOT_MINI_ complex acts processively in Smaug-dependent deadenylation. The substrate RNA in this assay was fluorescently labelled TCE^WT^-A_70_ RNA, which was used at 50 nM. The RNA was pre-incubated with or without 300 nM Smaug as indicated and the reaction started by the addition of ^Dm^CCR4-NOT_MINI_ (5 nM). Aliquots were withdrawn as indicated. Note that the time scale is in seconds. The asterisk indicates an unknown RNA species that we have not been able to remove. The graph on the right shows the accumulation of fully deadenylated product in the reaction containing Smaug.

To determine whether Smaug-dependent deadenylation is processive, we preincubated TCE^WT^ RNA (see Materials and Methods) with a threefold excess of Smaug over SREs. Limiting amounts of *^Dm^*CCR4-NOT_MINI_ were then added, and time-dependent deadenylation was measured. Biphasic kinetics were observed: In an initial burst phase, fully deadenylated end product was already present at the first time point, only five seconds after the start of the reaction, and continued to accumulate rapidly during the next ~ 20 seconds. At these early time points, most of the substrate RNA had not been attacked, and partially shortened intermediates of deadenylation were not visible. Thus, the reaction was processive. In the second phase of the reaction, additional deadenylation product accumulated at a much lower rate (**Fig. 6B**). Since the initial reaction was so fast, we could not test the prediction that the time at which the end product first appeared should be independent of enzyme concentration. However, the amount of RNA that was completely deadenylated within the burst phase was approximately stoichiometric with the CCR4-NOT complex: 5 nM CCR4-NOT complex deadenylated approximately 8 % of 50 nM substrate RNA within 5 s; 11% were completely deadenylated after 20 s. Thus, the initial burst phase represented the complete deadenylation of approximately one substrate RNA per enzyme complex, whereas the subsequent slow increase in deadenylated RNA presumably reflected the rate-limiting transition of the enzyme to new substrate molecules. In the control reaction lacking Smaug, no deadenylation was detectable under these conditions, but weak distributive activity was visible with longer incubation times (**Suppl. Fig. S4**).

### Cup also induces deadenylation

Additional constituents of the Smg-dependent repressor (Cup, Me31B, Tral, Belle) were also titrated individually into deadenylation assays containing the ^Dm^CCR4-NOT_FULL_ complex. Only Cup consistently stimulated deadenylation, also with the ^Dm^CCR4-NOT_MINI_ complex (**Fig. 7A; Suppl. Fig. 1C**). Thus, the Cup-dependent stimulation of deadenylation reported by (Igreja and Izaurralde, 2011) is a direct effect. The Cup-dependent reaction was also strongly stimulated by PEG (**Suppl. Fig. 2B**). Cup was separated into three non-overlapping fragments, the N-terminal, middle and C-terminal domains (N, M and C) (Igreja and Izaurralde, 2011). The three fragments as well as the NM and MC combinations were produced as His-λN-MBP fusion proteins (**Fig. 7B**). The phage λ N peptide (Baron-Benhamou et al., 2004) allowed binding to the deadenylation substrate, which contained two box B elements 16 nucleotides upstream of an A70 tail. All Cup fragments prepared in this manner were able to stimulate deadenylation (**Fig. 7C)**. No deadenylation was visible when Cup fragments were replaced by His-λN-MBP as a control, and the Cup fragments had no activity on their own. The ability of the M and C fragments to stimulate deadenylation is in agreement with (Igreja and Izaurralde, 2011). Surprisingly, all Cup fragments also stimulated deadenylation of a substrate RNA in which the box B elements were replaced by a control sequence (**Fig. 7C**); only the M fragment had weaker activity with this substrate.

**Figure 7.**
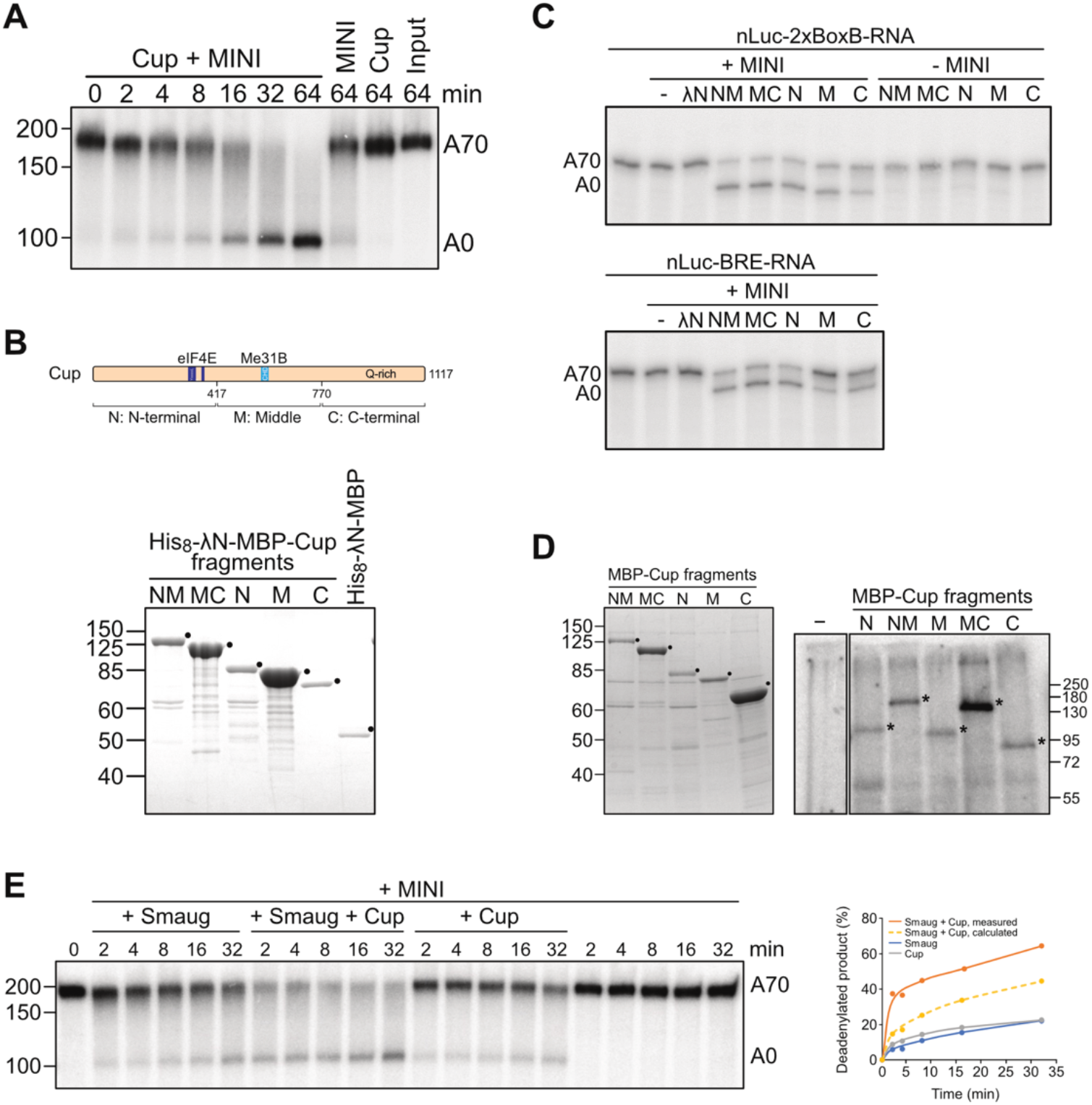
Cup also induces deadenylation. (A) Cup induces deadenylation by the ^Dm^CCR4-NOT_MINI_ complex. SRE^WT^-A_70_ RNA (20 nM) was preincubated with 80 nM Cup, and deadenylation was initiated by the addition of 2 nM ^Dm^CCR4-NOT_MINI_. Aliquots were withdrawn at different time points as indicated. Controls included incubations in the absence of Cup, with Cup only, or no protein added. (B) Preparation of Cup fragments. The top panel schematically shows the division of Cup into an N-terminal part (N) harbouring the eIF4E binding motifs, a middle part (M) with the Cup homology domain (CHD), and a C-terminal part (C) rich in glutamine residues. Bottom panel: SDS-polyacrylamide gel, stained with Coomassie, showing preparations of the three Cup fragments depicted in the top panel as well as the NM and MC combinations. All fragments had a His_8_-λN-MBP-tag. The λN peptide was utilized in tethering assays with BoxB-containing RNAs. The His_8_-λN-MBP protein served as a control. Desired proteins are labeled. (C) The ability of Cup to stimulate the CCR4-NOT complex is distributed over the protein. The His-λN-MBP-Cup fragments shown in (**B**) (40 nM each, except M, which was 80 nM) were pre-incubated for 15 min either with the nLuc-2xBoxB-A_70_ RNA (2 nM) or with the nLuc-BRE^MUT^-A_70_ RNA (2 nM). Deadenylation was started by the addition of ^Dm^CCR4-NOT_MINI_ (1 nM) and allowed to proceed for 30 min. Controls included incubations in the absence of CCR4-NOT, with CCR4-NOT only or with CCR4-NOT plus His_8_-λN-MBP. (D) Cup and all its fragments can be UV-crosslinked to RNA. Left panel: Coomassie-stained SDS gel showing MBP-Cup fragments. Right panel: Cup fragments shown in the left panel were UV-cross-linked to radiolabeled RNA. Cross-linking products were analyzed by SDS-polyacrylamide gel electrophoresis and autoradiography. (E) Smaug and Cup jointly stimulate deadenylation. Reactions were carried out in the absence of PEG. The SRE^WT^-A_70_ RNA (10 nM) was pre-incubated with Smaug (80 nM) or with buffer, then Cup (80 nM) or buffer was added for an additional 20 minutes. Deadenylation was started by the addition of ^Dm^CCR4-NOT_MINI_ (1 nM). Left panel: Analysis of reaction products by denaturing gel electrophoresis. Right panel: Completely deadenylated products were quantified. The broken yellow line indicates theoretical product accumulation predicted by additive behavior of Smaug and Cup.

The box B-independent activity of the Cup fragments and the activity of wild-type Cup suggest that the protein can bind RNA and recruit the CCR4-NOT complex. In fact, Cup has been identified as an RNA binding protein by proteome-wide UV cross-linking screens (Sysoev et al., 2016; Wessels et al., 2016), and so has the related protein 4E-T in mammals (Gerstberger et al., 2014). Unfortunately, RNA binding by purified Cup and its fragments could not be examined easily: The proteins did not yield interpretable results in electrophoretic mobility shift experiments and were not clean enough for nitrocellulose filter-binding experiments. Thus, UV cross-linking to a radiolabeled synthetic RNA oligonucleotide was used. For this purpose, all five Cup fragments were purified as MBP fusion proteins from baculovirus-infected Sf21 cells (**Fig. 7D**). All Cup fragments promoted deadenylation (**Suppl. Fig. 5A**), and all could be cross-linked to RNA (**Fig. 7D**). Migration of the major cross-link products in the SDS gel varied as expected from the molecular weights of the Cup variants, providing evidence that the signals were in fact derived from Cup. In order to further examine the identities of the cross-link products, we generated His-MBP fusions of full-length Cup (in Sf21 cells) and of the M, MC and C fragments (in *E. coli*). We were unable to generate the N and NM fragments with this type of fusion. Fusion proteins were cross-linked to RNA, and their identities were ascertained by their binding, under denaturing conditions, to a Ni-NTA matrix (**Suppl. Fig. 5B**). Some proteolytic products common to the M and MC fragments were copurified on the Ni column, but a contamination common to all fragments was not, confirming the specificity of the pull-down. Thus, full-length Cup as well as all fragments tested are able to bind RNA.

For the purpose of testing whether Smg and Cup can simultaneously stimulate CCR4-NOT-dependent deadenylation, the reaction was weakened by omission of PEG from the buffer. Under these sensitized conditions the simultaneous presence of Smaug and Cup indeed led to significantly improved deadenylation of the SRE^WT^ RNA compared to either protein alone; stimulatory effects were reproducibly more than additive (**Fig. 7E**). Each protein was used at a concentration that individually was saturating for deadenylation and close to saturation in RNA binding **(Suppl. Fig. 5C, D)**. Thus, Smg and Cup bound to the same RNA can cooperate in the stimulation of the CCR4-NOT complex.

## DISCUSSION

During early embryonic development of *Drosophila,* Smaug is responsible for the degradation of hundreds of maternal mRNAs, thus making a major contribution to the maternal-to-zygotic transition (Chen et al., 2014a; Tadros et al., 2007). Its best-studied target is the *nanos* (*nos*) mRNA. Smaug represses *nos* both by preventing its translation and by inducing its deadenylation by the CCR4-NOT complex. As Smaug binds the *nos* 3’-UTR in the company of six other proteins, the mechanism by which it induces deadenylation is potentially complex. In order to examine the mechanism of the deadenylation reaction, we have reconstituted the eight subunit *Drosophila* CCR4-NOT complex and also overproduced and purified Smaug as well as the other constituents of the Smaug-dependent repressor complex. Activity assays with these proteins revealed that Smaug on its own is able to induce an efficient, processive deadenylation by the CCR4-NOT complex. As a second component of the repressor complex, Cup was also able to stimulate CCR4-NOT-catalyzed deadenylation. A third component, PABPC, modestly promoted deadenylation, although this is unlikely to contribute specifically to *nos* repression as PABPC is thought to bind the poly(A) tails of all mRNAs. The other components of the repressor complex did not facilitate deadenylation in our *in vitro* assays. The inactive components included Me31B, even though the protein is able to bind NOT1 (Chen et al., 2014b; Mathys et al., 2014; Rouya et al., 2014).

Our data support a simple tethering model for the ability of Smaug to accelerate deadenylation: Smaug binds SREs with high affinity (Aviv et al., 2003; Green et al., 2003). As shown here, the protein also interacts directly with the CCR4-NOT complex via the C-terminal domain of NOT3 and induces a processive activity of CCR4-NOT. Smaug-induced deadenylation proceeded at a rate too fast to be measured by manual pipetting. The amount of RNA deadenylated in the initial burst also appeared to be stoichiometric with respect to the deadenylase, although this conclusion is limited by the accuracy with which the concentration of CCR4-NOT could be determined. These data indicate that Smaug promotes deadenylation by preventing the dissociation of the deadenylase from its substrate. This corresponds to the mechanism by which deadenylation effectors Mmi1, Puf3 and Zfs1 from *S. pombe* stimulate their cognate CCR4-NOT complex (Stowell et al., 2016; Webster et al., 2019). Smaug-dependent deadenylation was initially speculated to be more complex, as the reaction appeared to be ATP-dependent in extracts of S2 cells (Jeske et al., 2006). Deadenylation induced by miRNAs behaved similarly (Iwasaki et al., 2009). However, the apparent ATP dependence was later explained by the accumulation of AMP upon ATP depletion; AMP inhibited deadenylation (Niinuma and Tomari, 2017). We have independently found that the apparent ATP-dependence of deadenylation in nuclear extract was only observed with substrate RNAs produced by run-off transcription that ended in a few non-A residues downstream of the poly(A) tail due to the restriction site used at the time to linearize the template; deadenylation was not inhibited by ATP depletion when the poly(A) tails were added by poly(A) polymerase (C. Temme and EW, unpublished data). Perhaps degradation of the non-A residues is more sensitive to AMP inhibition. Regardless, our reconstitution experiments confirm that Smaug-dependent deadenylation does not require ATP.

Deadenylation effectors typically interact with CCR4-NOT in a complex manner, employing multiple short interaction motifs embedded in intrinsically disordered regions (Bhandari et al., 2014; Stowell et al., 2016; Webster et al., 2019). For example, contacts to CCR4-NOT involve four domains of the *Drosophila* Pumilio protein (Arvola et al., 2020) or multiple tryptophan-containing motifs of the miRNA-associated GW182 proteins (Chekulaeva et al., 2011; Chen et al., 2014b; Mathys et al., 2014). Many effectors also interact with more than one subunit of CCR4-NOT. For example, the *Drosophila* Nanos protein is itself an effector of deadenylation and uses redundant binding sites to recruit CCR4-NOT: The dominant site forms one short α-helix contacting NOT1 in its C-terminal domain and a second α-helix binding the NOT box of NOT3 (Raisch et al., 2016). Likewise, *Drosophila* Roquin has a binding site for CAF40 and redundant binding sites for the NOT module (Sgromo et al., 2017), and mammalian TTP interacts both with NOT1 (Fabian et al., 2013) and CAF40 (Bulbrook et al., 2018). In the case of Smaug, the C-terminal region of NOT3 provides the major interaction surface. However, multiple regions of Smaug appear to be involved as we were unable to map an individual NOT3 binding domain in the protein.

In addition to Smaug, Cup was also able, on its own, to stimulate CCR4-NOT-dependent deadenylation. The ability to induce deadenylation seems to be distributed over much of the protein, as each of the three non-overlapping fragments was active. Our results are consistent with the report (Igreja and Izaurralde, 2011) that the M and C fragments of Cup associate with CCR4-NOT and, when tethered to a reporter RNA, induce deadenylation in cells. Our results in the reconstituted *in vitro* system indicate that this effect of Cup is direct. In spite of very limited sequence similarity (Kamenska et al., 2014), the Cup-related protein 4E-T also interacts with CCR4-NOT and induces deadenylation in tethering assays (Räsch et al., 2020). The ability of Cup and its fragments to induce deadenylation even in the absence of tethering suggests they are able to bind RNA. Unfortunately, we have been unable to purify Cup or its fragments to homogeneity, as the proteins did not elute in clean peaks from any column tested. Thus, RNA binding activity in nitrocellulose filter-binding assays could not be attributed to Cup as opposed to contaminations. Upon electrophoresis in native gels, all complexes formed between RNA and the Cup preparation remained stuck in the wells so that, again, the protein responsible for binding could not be identified with certainty. However, UV cross-linking assays were consistent with RNA binding of Cup and its fragments.

Although Cup is able to bind RNA and induce deadenylation on its own, its *in vivo* activity probably depends on the protein being recruited to specific mRNAs by Smaug and other RNA binding proteins, for example Bruno (Chekulaeva et al., 2006; Nakamura et al., 2004). The presence of two deadenylation effectors, Smaug and Cup, in a single complex is not without precedent: Nanos and Pumilio and, in some cases, Brat, cooperate in the regulation of *Drosophila hunchback* and other mRNAs (Arvola et al., 2017; Joly et al., 2013). Both Nanos and Pumilio can individually elicit deadenylation by CCR4-NOT (see above) (Arvola et al., 2020; Raisch et al., 2016; Van Etten et al., 2012). When both Smaug and Cup were used at near-saturating concentrations, their effects on deadenylation were not merely additive, but a modest degree of cooperativity was reproducible observed. This cooperative effect was seen in the absence of a crowding reagent; in the presence of PEG, the Smaug-dependent reaction is so efficient that one cannot expect an additional stimulation by Cup.

Previous knock-down experiments and overexpression of catalytically dead polypeptides in cultured *Drosophila* cells suggested that CAF1 may be the dominant catalytic subunit of CCR4-NOT *in vivo* (Arvola et al., 2020; Temme et al., 2010). The interpretation of these experiments is limited by incomplete protein depletion and an uncertain degree to which WT protein is replaced by the inactive version in the CCR4-NOT complex. CCR4 is important for the deadenylation of *nos* (Zaessinger et al., 2006) (see below), and genetic evidence for the importance of CCR4 catalytic activity has been presented (Joly et al., 2013). Our reconstitution experiments now provide clear evidence that both CCR4 and CAF1 are active nucleases. Individual inactivation of either subunit reduced the activity of the complex by more than 50%. This apparent interdependence of the two activities may be most easily explained by the inactive subunit transiently blocking access to the 3’ end. Nevertheless, inactivation of either CCR4 or CAF1 reduced the activity of the complex to a similar extent, both in unassisted, basal deadenylation and in the Smaug-dependent reaction, suggesting comparable contributions of the two nucleases to the activity of CCR4-NOT. However, since the activity of CAF1 is highly sensitive to pH and the concentrations of Mg^2+^ and Zn^2+^ (Chen et al., 2021), our results do not exclude the possibility that one catalytic subunit plays a dominant role *in vivo*.

Using a fully reconstituted CCR4-NOT complex from *S. pombe,* (Webster et al., 2018) found that the enzyme’s activity is facilitated by PABPC (Pab1 in *S. pombe*). Similar conclusions have been reached for the human deadenylase (Yi et al., 2018), although reconstitution was limited to a CCR4-CAF1 heterodimer. In accordance with these data, we also observed a modest stimulation of the *Drosophila* CCR4-NOT complex by PABPC. When added to a Smaug-containing deadenylation reaction, which is highly efficient to begin with, PABPC caused a weak additional stimulation. In contrast, (Webster et al., 2019) reported a weak inhibitory effect of PABPC on the activity of *S. pombe* CCR4-NOT activated by an RNA binding protein. In summary, it is probably fair to say that the effects of the deadenylation effectors that have been tested are dominant over the effects of PABPC.

(Webster et al., 2018) and (Yi et al., 2018) found that the catalytic activity of *S. pombe* or human CCR4 is stimulated by PABPC, whereas CAF1 is unable to degrade PABPC-bound poly(A). In contrast, we observed that both catalytic subunits of *Drosophila* CCR4-NOT are able to cope with a PABPC-poly(A) complex. This discrepancy may reflect species-specific differences. However, (Stupfler et al., 2016) reported that mouse CAF1 (CNOT7) digested PABPC-bound poly(A) and the naked polymer with similar efficiencies; thus, it seems more likely that the effect of PABPC depends on the reaction conditions, to which CAF1 is highly sensitive. Mice lacking both CCR4 orthologues (CNOT6 and CNOT6L) are viable, have no overt phenotype, and fibroblasts derived from them have a normal poly(A) tail length distribution. In contrast, the deletion of NOT1 or combined deletion of both CAF1 orthologues (CNOT7 and CNOT8) are lethal, and their reduced expression leads to an accumulation of long poly(A) tails (Mostafa et al., 2020). These data show that CCR4-NOT is essential, but CCR4-type subunits are not. Thus, CAF1-type subunits must be able to degrade PABPC-covered poly(A) in vivo; any impediment imposed by PABPC cannot be absolute. The ability of CAF1 to degrade poly(A) in the presence of PABPC is also enhanced in the presence of BTG2/TOB (Ezzeddine et al., 2007; Pavanello et al., 2018; Stupfler et al., 2016).

Our data reveal that Cup plays a direct role in the deadenylation of *nos* and probably many other RNAs. Smaug, acting on CCR4-NOT by itself and promoting the recruitment of Cup (Nelson et al., 2004), accelerates deadenylation directly and presumably also indirectly. However, the experiments capture only select aspects of the complicated *in vivo* situation: Smaug and its partner Cup are not the only effectors of *nos* deadenylation; piRNAs are also involved (Rouget et al., 2010), and accordingly a deletion of Smaug only partially stabilizes *nos* RNA (Semotok and Lipshitz, 2007). Whereas *nos* loses its entire poly(A) tail in the Smaug- and CCR4-NOT-dependent *in vitro* reaction, the RNA persists with an oligo(A) tail *in vivo* (see Introduction). Possibly, the complete and extremely rapid deadenylation reaction observed *in vitro* is tempered by conditions prevailing in the embryo. However, a competing poly(A) tail extension reaction catalyzed by the non-canoncial poly(A) polymerase GLD2 (encoded by *wispy*) presumably plays a more important role (Benoit et al., 2008).

In the early *Drosophila* embryo, translational efficiency is tied to poly(A) tail length (Eichhorn et al., 2016). Therefore, Smaug-induced deadenylation, even though it does not go to completion in the embryo, contributes to the translational repression of the non-localized fraction of *nos* RNA (Zaessinger et al., 2006). However, *nos* translation is also repressed by mechanisms independent of deadenylation (Götze et al., 2017; Jeske et al., 2011; Nelson et al., 2004). Although CCR4-NOT can repress translation independently of its deadenylation activity (Braun et al., 2011; Chekulaeva et al., 2011; Cooke et al., 2010; Waghray et al., 2015), the deadenylase is an unstable and/or substoichiometric constituent of the Smaug-dependent repressor complex (Götze et al., 2017) and therefore unlikely to play a major role in repression beyond its deadenylating activity. Deadenylation not only contributes to translational repression, but is also the first step in the decay of *nos*, as shown by the stabilization of *nos* in *smg* and *twin* mutants (Zaessinger et al., 2006). Cup stabilizes deadenylated RNA by inhibiting decapping (Igreja and Izaurralde, 2011). Thus, the degradation of Cup during the first three hours of embryonic development (Cao et al., 2020) might be thought to be essential for the further degradation of deadenylated *nos*. However, *nos* is cleared rapidly and completely in *wispy* mutants (Benoit et al., 2008); thus, even in the presence of Cup, deadenylation appears to be sufficient to induce the decay of the *nos* mRNA.

## MATERIALS AND METHODS

### Expression clones and viruses

Co-expression of two or more proteins from one baculovirus vector was performed by means of the MultiBac expression system (Berger et al., 2004; Fitzgerald et al., 2006). For expression of the ^Dm^CCR4-NOT variants, all ORFs were amplified from *D. melanogaster* cDNA. For NOT1_MINI_ (amino acid residues1147-2505), NOT2, NOT3 and Caf1 an N-terminal His_8_-tag was added by PCR. A C-terminal FLAG-tag was appended to CCR4 and CAF40 by PCR. His_8_-NOT2 was cloned into the pFBDM vector between the SmaI and KpnI restriction sites. His_8_-NOT1_MINI_ (residues 1147-2505) was first introduced between the NheI and HindIII restriction sites of the pET28a-MBP plasmid, then cut out with PspOMI and HindIII and inserted into the pFBDM-His_8_-NOT2 plasmid between the NotI and HindIII restriction sites. This pFBDM-His_8_-NOT2_His_8_-NOT1_MINI_ plasmid was the basis for the generation of longer NOT1 variants lacking the His tag: Additional parts of NOT1 were inserted into the NOT1_MINI_ plasmid between the XbaI and AgeI restriction sites until the full length NOT1 was obtained: AvrII_NOT1_MINI_EXT__AgeI (residues 878-1570); XbaI_NOT1_pE__AgeI (residues 288-1570); XbaI_NOT1_pC__AgeI (residues 1-1570). Caf40-FLAG was cloned into the pFBDM vector between the EcoRI and HindIII restriction sites. His_8_-NOT3 was then inserted between the SmaI and KpnI restriction sites of pFBDM-Caf40-FLAG, resulting in the pFBDM-Caf40-FLAG_His_8_-Not3 plasmid. For the nuclease module, CCR4-FLAG was inserted in the pFBDM vector between the XhoI and KpnI restriction sites. His_8_-Caf1 was then cloned between the BamHI and XbaI restriction sites of pFBDM-CCR4-FLAG, resulting in the pFBDM-CCR4-FLAG_His_8_-Caf1 plasmid. Point mutations in CCR4 (412D/414N to alanine) and Caf1 (53D/55E to alanine) (Temme et al., 2010) were introduced by overlap extension PCR. NOT10 and NOT11 cDNAs, cloned in pnYC-NpM and pnEA-CvH, respectively (Bawankar et al., 2013; Diebold et al., 2011) were obtained from Eugene Valkov. The NOT10 sequence was removed from its vector with KpnI and NheI and inserted between the KpnI and AvrII sites of pnEK-His_8_-MBP-Cup (described below), replacing the Cup ORF. Thereby, an N-terminal His_8_-MBP-tag followed by a HRV 3C protease site was fused to NOT10. A BglII-XbaI fragment containing His_8_-MBP-NOT10 was then cut out of the pnEK vector and cloned between the BglII and AvrII restriction sites behind the second T7 promoter of the pETDuet-1 vector (Merck, Darmstadt, Germany). The resulting plasmid was cut with SalI and BglII, and NOT11, cut as an XhoI-BamHI fragment from its vector, was introduced behind the first T7 promoter. This created a fusion with an N-terminal His_6_-tag in the pETDuet-1-His_8_-MBP-NOT10_His_6_-NOT11 plasmid.

DNA sequences coding for the putative constituents of the SRE-dependent repressor complex were also amplified from *D. melanogaster* cDNA, except Cup, which was a kind gift of Fulvia Bono (in the pnEK vector (Busso et al., 2011)). C-terminal FLAG-tags were added to Smaug and Cup by PCR. Smaug-FLAG was inserted between the NheI and NotI restriction sites of the pFBDM vector. Truncated Smaug variants were amplified from pFBDM-Smaug-FLAG. Smaug N (aa 1-329) was inserted between the BamHI and SmaI restriction sites, Smaug MC (aa 585-999) between the EcoRI and BsaAI restriction sites and Smaug C (aa 661-999) between the EcoRI and SmaI restriction sites of the pGEX6p1 vector. Smaug NM (aa1-660) was inserted between the BamHI and Eco53kI restriction sites of the pET28a vector. Afterwards, Smaug NM was cut from its vector with BamHI and XhoI and introduced between the same restriction sites into the pET28a-His_8_-MBP plasmid, thereby receiving an N-terminal His_8_-MBP-tag. His_8_-MBP-SmaugNM was cut out of the pET28a vector with XbaI and XhoI and inserted into the pFastBac1 vector between the SpeI and XhoI restriction sites. A Smaug M (aa 587-851) clone, inserted between the BsaI and XhoI restriction sites of the pET-SUMOadapt vector (Bosse-Doenecke et al., 2008), was generated by Bodo Moritz. Cup-FLAG was cloned between the SmaI and XhoI restriction sites of the pFBDM vector. For the generation of MBP-tagged Cup fragments, MBP was cut out of pnYC-NpM (Diebold et al., 2011) with SpeI and XhoI and inserted between the equivalent sites of pnEK-Cup. An N-terminal His_8_ tag was cloned, as a synthetic oligonucleotide, into the NcoI site of pnEK-MBP-Cup. The resulting plasmid, pnEK-His_8_-MBP-Cup, was the basis for the generation of truncated Cup variants (N: aa 1-417; M: aa 418-770; C: aa 771-1117; and combinations thereof), which were amplified from pFBDM-Cup-FLAG and inserted between the XhoI and AvrII restriction sites of pnEK-His_8_-MBP-Cup for *E. coli* expression. MBP-Cup fragments were cut out of the pnEK-His_8_-MBP vector with NcoI and AvrII and inserted into the pFBDM vector between the NcoI and NheI restriction sites. An N-terminal His_8_-λN-tag was introduced as a restriction fragment into the pFBDM-MBP-Cup fragments using the NcoI and SmaI sites, resulting in the pFBDM-His_8_-λN-MBP-Cup fragment plasmids. The GST-Me31B clone in pFastBac1 (Thermo Fisher) has been described (Götze et al., 2017). A C-terminal His_8_-tag was appended to Tral by PCR, and the resulting fragment was cloned into pFastBac 1 with EcoRI and NcoI. For expression of His-T7-Tral-His in *E. coli,* the coding sequence was cut out from the pFastBac 1 construct and transferred to pET28a (Merck, Darmstadt, Germany) by means of HindIII and Eco53kI. An N-terminal His_8_-tag was added to Belle by PCR, and His_8_-Belle was inserted between the XhoI and HindIII restriction sites of the pFastBac1 vector. PABPC was cloned into pET-28a using BamHI and XhoI.

All expression constructs were confirmed by Sanger sequencing. All primers used to generate expression constructs are listed in **Supplemental Table 1.**

pFastBac1 and pFBDM DNA constructs were transformed into DH10MultiBac cells (Geneva Biotech) and incubated for 5 hours to allow transposition into the mini-*att*Tn7 sites of the bacmid DNA. Colonies were selected for correct bacmid DNA by antibiotic resistance and blue-white screening. Bacmid DNA was isolated via alkaline lysis (Qiagen buffers P1, P2 and P3) and isopropanol precipitation. Sf21 cells (1.5×10^6^ cells seeded into a 6-well plate) were transfected with 10 μg bacmid DNA, which had been pre-incubated in 200 μL Ex-Cell 420 medium plus 5 μL FuGENE HD transfection reagent (Promega) for 20 minutes at 27 °C. After 96 hours, the supernatant was collected and used to infect 0.8 × 10^6^ Sf21 cells/ml at a 1:500 volume ratio for virus propagation (V1 generation). The V1 generation was used to infect Sf21 cells for protein expression.

### Protein expression and purification

Anti-FLAG M2 agarose and FLAG peptide were from Sigma, Ni-NTA agarose was from Qiagen, amylose resin from NEB, and Glutathione Sepharose 4B and the Superose 6 column from Cytiva. PES concentrators were from Thermo Fisher and Amicon concentrators from Merck, Darmstadt, Germany. HRV 3C protease was purified in-house by Bodo Moritz. Sf21 cells were grown as suspension cultures in Ex-Cell 420 serum-free medium (Sigma-Aldrich) at 27 °C.

For the reconstitution of hexameric ^Dm^CCR4-NOT complexes, Sf21 cells were coinfected with a baculovirus expressing His_8_-NOT3 + Caf40-FLAG, a second virus expressing CCR4-FLAG + His_8_-Caf1, and a third virus expressing His_8_-NOT2 with one of the different NOT1 variants. The baculoviruses were used at a 2:1:2 ratio. Cells were harvested 72 hours after infection, resuspended in ice-cold lysis buffer (500 mM NaCl, 50 mM HEPES-NaOH and 10 mM potassium phosphate, pH 7.6, 10 % sucrose, 1 mM PMSF, 1 μM pepstatin A). Sucrose stabilized the CCR4-NOT complex during freezing and thawing. Cells were lysed by sonication on ice, the lysate was cleared by centrifugation for 20 min at 20.000 x g and the supernatant applied to anti-FLAG M2 agarose matrix for 2 hours under constant rotation at 6 to 8 °C. The matrix was washed four times with wash buffer (lysis buffer minus PMSF and pepstatin A) in a batch format, then protein was eluted by addition of wash buffer with 200 μg/ml FLAG peptide.

The eluate was collected after 30 min, concentrated with a PES concentrator (10 kDa MWCO) and applied to a Superose 6 column equilibrated in wash buffer. Removal of the filter from the Superose column improved protein recovery. Fractions containing the hexameric complex were pooled, concentrated as before and frozen in liquid nitrogen before final storage at −80 °C.

For reconstitution of the octameric ^Dm^CCR4-NOT_FULL_ complex, His_8_-MBP-NOT10 and His_6_-NOT11 were co-expressed in *E. coli* BL21 Star (DE3) cells with 24 h lactose autoinduction (Studier, 2005). Cells were harvested, resuspended in ice-cold lysis buffer containing 1 mg/ml lysozyme and 10 mM imidazole and incubated for 1 hour under constant rotation at 6 to 8 °C. Cells were then sonicated, the lysate was centrifuged for 20 min at 15.000 x g and the supernatant applied to Ni-NTA agarose for 2 h under constant rotation at 6 to 8 °C. The matrix was washed four times with wash buffer plus 20 mM imidazole, and proteins were eluted with wash buffer plus 160 mM imidazole. After 30 min, the eluate was applied to amylose resin equilibrated in wash buffer and incubated for 2 hours. NOT10 and His_6_-NOT11 were eluted by a 2 h incubation with an approximately equimolar amount of HRV 3C protease and concentrated with a PES concentrator (10 kDa MWCO). A twofold excess of the heterodimer was incubated for 2 h at 8 °C with the hexameric CCR4-NOT (isoform NOT1_PC_) to reconstitute ^Dm^CCR4-NOT_FULL_, which was purified on a Superose 6 column, concentrated and stored like the hexameric complex.

For production of Smaug-FLAG, Cup-FLAG, Tral-His_8_, GST-Me31B or His_8_-Belle, Sf21 cells were infected with the relevant baculoviruses. Infected cells were processed as above up to the addition of the affinity matrix. Binding to the respective matrix (anti-FLAG M2 agarose, Ni-NTA agarose, or Glutathione Sepharose 4B) was carried out for 2 h, then the matrix was washed four times with wash buffer (with 20 mM imidazole for His_8_-tagged constructs), and proteins were eluted by wash buffer containing 200 μg/ml FLAG peptide or 160 mM imidazole or by on-column cleavage with HRV 3C protease for Me31B. After elution, the proteins were, if necessary, concentrated with an Amicon Ultra concentrator (10 kDa MWCO), flash frozen and stored at −80 °C. MBP- and His_8_-λN-MBP-Cup fragments and His_8_-MBP-SmaugNM were also expressed in Sf21 cells by baculoviral infection. Infected cells were processed as above up to the addition of the affinity matrix. His-MBP-tagged proteins were first purified by Ni-NTA affinity chromatography as above and then applied for 2 h to an amylose agarose matrix (NEB). Bound proteins were eluted with wash buffer containing 20 mM maltose. MBP-Cup fragments were purified by direct addition of cell lysates to the amylose matrix. The remaining Smaug fragments (MC, N, M and C) were expressed in *E. coli* BL21 Star (DE3) and processed as described for NOT10-NOT11 up to the addition of the affinity matrix. His_6_-SUMO-SmaugM was purified by Ni-NTA affinity chromatography like NOT10-NOT11. The other fragments were bound to Glutathione Sepharose 4B and eluted with wash buffer plus 20 mM glutathione.

*E. coli* BL21 Star (DE3) cells were used for the expression of His_6_-T7-PABPC and His_6_-T7-Tral-His_8_. Expression and Ni-NTA chromatography were carried out as described for the NOT10-NOT11 purification, except that the buffer contained no phosphate. At this point, Tral and PABPC were flash frozen and stored at −80°C.

Identities of purified proteins were confirmed by western blotting. Protein concentrations were determined by SDS-polyacrylamide gel electrophoresis and Coomassie staining in comparison to a set of BSA standards. Intensities of protein bands were evaluated with Fiji. For deadenylation experiments, proteins were pre-diluted in 200 mM potassium acetate, 50 mM HEPES-KOH, pH 7.4.

### Western blotting

Proteins were transferred onto nitrocellulose membrane (GE Healthcare) by wet blotting overnight at 8°C and 27 V. Membranes were blocked with 1.5 % gelatin (from cold water fish skin; Sigma-Aldrich) in 1x TBS - 0.05 %Tween and incubated with the primary antibody for 2 h. The antibody against Smaug (Chartier et al., 2015) was diluted 1:500 in blocking buffer. The membrane was washed three times with 1x TBS-Tween and incubated with fluorescently labelled secondary antibody (IR-Dye, LI-COR; 1:15,000). The membrane was washed again three times with 1x TBS-Tween and scanned with a LI-COR Odyssey imager.

### Deadenylation substrates

The SRE-only RNA and the TCE RNA as well as their mutant (SRE^MUT^) versions have been described (Jeske et al., 2006). The TCE RNA was called *nos* RNA in the Jeske et al. paper. The plasmid vector for these RNAs also encoded a poly(A) tail of about 70 nt. Compared to the version described (Jeske et al., 2006), it was modified by introduction of a BsaI site into the HindIII site at the end of the poly(A) tail, so that, upon BsaI digestion, a run-off transcript was produced that ended in a straight poly(A) tail without additional nucleotides from the restriction site. The BoxB tethering construct was generated by assembly of a synthetic BamHI-XbaI restriction fragment containing two BoxB elements (Baron-Benhamou et al., 2004) (**Supplemental Table 1**) and insertion into the BglII-XbaI restriction sites of the pBSK-nLuc-nos plasmid (Kluge et al., 2020). The control construct (nLuc-BRE^MUT^-A_70_ RNA) contained the mutated AB Bruno recognition element (BRE) of the *oskar* 3’ UTR (Kim-Ha et al., 1995). The pBSK-nLuc-BRE^WT^ plasmid was first generated by amplifying the BRE from *Drosophila* cDNA and introducing it between the BglII-XbaI restriction sites of the pBSK-nLuc-nos plasmid. The mutant BRE was then assembled from four synthetic DNA oligonucleotides (**Supplemental Table 1**) and introduced downstream of the nLuc ORF between the BamHI and EcoRI sites of the pBSK-nLuc-BRE^WT^ plasmid, replacing the WT BRE.

RNAs were synthesized with T3 RNA polymerase (Promega) in the presence of [α-^32^P]UTP and 7 mM “anti-reverse” cap analog (m^7,3’-O^GpppG; Jena Bioscience). All radiolabeled RNAs were gel-purified. Alternatively, the RNA was labeled by incorporation of a 6-carboxyfluorescein-(6-FAM-) labelled ApG cap analog. The cap analog was added to the transcription reaction at 1 mM in the absence of GTP. After a 5 min preincubation, GTP was added to 0.2 mM, and the incubation was continued for an additional 20 minutes before the GTP concentration was raised to 1 mM. After an additional hour, the reaction was incubated with 1 U DNase I (Roche), the RNA was phenol-extracted and ethanol precipitated and used without further purification.

The chemically synthesized FAM 7mer-A_20_ RNA has been described (Raisch et al., 2019).

### Deadenylation assays

The reaction buffer consisted of 50 mM potassium acetate, 30 mM HEPES-KOH, pH 7.4, 2 mM magnesium acetate, 0.15 mg/mL nuclease-free BSA (Merck-Millipore), 2 mM DTT, 800 U/mL murine RNase Inhibitor (NEB) and 3 % (w/v) PEG 20,000. The presence of BSA in the buffer improved the stability of the CCR4-NOT complex during extended incubations at low concentration. Yeast tRNA (0.25 mg/mL) was added as a carrier. Reactions with the FAM-7mer-A_20_ RNA did not contain tRNA, unless noted otherwise. Substrate RNA was either directly incubated with CCR4-NOT complexes or pre-incubated for 20 - 30 min with potential activators of deadenylation at 25 °C, and deadenylation was started by addition of CCR4-NOT. Reaction temperature was 25 °C both for the human and the *Drosophila* CCR4-NOT complex. The reaction was stopped by addition of a two-to threefold excess of ice-cold formamide loading buffer (95% deionized formamide, 17.5 mM EDTA, pH 8.0, 0.01% bromophenol blue, 0.01% xylene cyanol). Xylene cyanol was omitted for FAM-7mer-A_20_ substrates. Samples were heated to 95 °C for 3 min, cooled on ice and separated on denaturing polyacrylamide gels (19:1 acrylamide-bis acrylamide, 1x TBE, 7 M urea). Fluorescent reaction products were directly visualized by scanning with a Typhoon 9200, whereas gels with radioactive RNA were placed on storage phosphor screens overnight at −20 °C and the screens scanned the next day. Images were analysed using Fiji (Schindelin et al., 2012).

Deadenylation assays and other experiments were performed at least twice, and representative experiments are shown.

### RNA binding assays

His_8_-MBP- or MBP-tagged Cup or Cup fragments were incubated with a 5’-^32^P-labeled RNA oligonucleotide (GGGTTTAGTGCGCACGTG, 18nt) in a reaction mixture containing 16 mM HEPES, pH 7.6, 50 mM potassium acetate, 1 mM magnesium acetate, 0.8 mM ATP, 0.24 mg/ml yeast tRNA, 1 mM DTT, 10 % glycerol for 15 min at room temperature and UV crosslinked (Stratalinker 1800, 254 nm, 200 mJ/cm^2^) on ice. Reaction products were separated on SDS-polyacrylamide gels, which were dried and analyzed by phosphoimaging. For analysis of the cross-link product by affinity purification, a part of each reaction was stored as input. The remaining portions were diluted in 50 mM HEPES, pH 7.6, 500 mM KCl, 10 % sucrose, 6 M urea, 0.05 % NP40 and incubated for 15 min at room temperature. Magnetic Ni-NTA beads (Promega) were added to the reaction, and the mix was incubated at room temperature for 1 h on a rotating wheel. The matrix was collected and washed twice with buffer as above. Before elution in SDS sample buffer, the matrix was transferred to a fresh tube. Input and elution fractions were analyzed by gel electrophoresis as above.

For electrophoretic mobility shift assays, binding reactions were carried out in deadenylation reaction buffer, including tRNA, plus 5 % glycerol for 20 to 30 minutes at 25 °C. RNA-protein complexes were separated on nondenaturing polyacrylamide gels (5% 60:1 acrylamide-bis acrylamide, 0.5x TBE) and visualized by phosphoimaging.

### Protein-protein interaction assays

ReLo assays were performed as described (Salgania et al., 2022). In short, *Drosophila* S2R+ cells were seeded onto four-well chambered coverslips (Ibidi), co-transfected with the desired combination of pAc5.1 plasmids expressing the two proteins of interest, and protein localization was analyzed after two days by live confocal fluorescence microscopy. Split-ubiquitin yeast two-hybrid experiments were performed as described (Jeske et al., 2015). The desired combinations of bait and prey plasmids were co-transformed into NMY51 cells and plated onto SC agar lacking Leu and Trp. For the spotting assay, several colonies were pooled to prepare an overnight liquid culture. Three 10-fold dilutions were spotted onto SC agar plates either lacking Trp and Leu (control) or Trp, Leu, and His (selection), and, after three days of growth, images were taken. Detailed information on all plasmids used for protein-protein interaction assays is provided in **Supplemental Table 2**.

To test for an interaction between Smaug and the CCR4-NOT complex by gel filtration, Smaug (15 μg in 50 μL) was incubated either with ^Dm^CCR4-NOT_MINI_ (15 μg in 20 μL) or protein buffer for 1 h at 8 °C. 50 μL of the mixture was applied to a Superose 6 gel filtration column (bed volume ~ 2.4 mL; Cytiva) equilibrated in lysis buffer lacking PMSF and pepstatin A. The column was run with the same buffer at 0.017 column volumes per min. Column fractions were analyzed by SDS polyacrylamide gel electrophoresis and Coomassie staining and/or by western blotting with an antibody against Smaug.

## ACKNOWLEDGEMENTS

We are grateful to Eugene Valkov and Yevgen Levdansky for generously providing the purified human CCR4-NOT samples, cDNAs for *Drosophila* NOT10 and NOT11 and FAM-7mer-A_20_ RNA and for helpful discussions and comments on an earlier version of the manuscript; to Fulvia Bono for a Cup cDNA clone; to Larissa Möckel for help with RNA binding assays; to Bodo Moritz for HRV 3C protease and the Smaug M clone; to Maik Schauerte, Felia Haffelder and Corinna Nowak for generating plasmids used for interaction studies. We thank the Nikon Imaging Center at the University of Heidelberg for access to microscopes and gratefully acknowledge the data storage service SDS@hd supported by the Ministry of Science, Research and the Arts Baden-Württemberg (MWK) and the German Research Foundation (DFG) through grants INST 35/1314-1 FUGG and INST 35/1503-1 FUGG. This work was supported by DFG grants to EW (WA 548/16-1 and WA 548/17-1 within the framework of SPP 1935) and by an Emmy Noether grant of the DFG to MJ (JE-827/1-1).

## REFERENCES

Alberti, S., Gladfelter, A., and Mittag, T. (2019). Considerations and Challenges in Studying Liquid-Liquid Phase Separation and Biomolecular Condensates. Cell 176, 419–434.

Alles, J., Legnini, I., Pacelli, M., and Rajewsky, N. (2021). Rapid nuclear deadenylation of mammalian messenger RNA. Biorxiv, https://doi.org/10.1101/2021.1111.1116.468655

Arvola, R.M., Chang, C.T., Buytendorp, J.P., Levdansky, Y., Valkov, E., Freddolino, P.L., and Goldstrohm, A.C. (2020). Unique repression domains of Pumilio utilize deadenylation and decapping factors to accelerate destruction of target mRNAs. Nucleic Acids Res 48, 1843–1871.

Arvola, R.M., Weidmann, C.A., Tanaka Hall, T.M., and Goldstrohm, A.C. (2017). Combinatorial control of messenger RNAs by Pumilio, Nanos and Brain Tumor Proteins. RNA Biol 14, 1445–1456.

Aviv, T., Lin, Z., Lau, S., Rendl, L.M., Sicheri, F., and Smibert, C.A. (2003). The RNA binding SAM domain of Smaug defines a new family of post-transcriptional regulators. Nature Struct Biol 10, 614–621.

Baron-Benhamou, J., Gehring, N.H., Kulozik, A.E., and Hentze, M.W. (2004). Using the lambdaN peptide to tether proteins to RNAs. Methods Mol Biol 257, 135–154.

Bashirullah, A., Halsell, S.R., Cooperstock, R.L., Kloc, M., Karaiskakis, A., Fisher, W.W., Fu, W., Hamilton, J.K., Etkin, L.D., and Lipshitz, H.D. (1999). Joint action of two RNA degradation pathways controls the timing of maternal transcript elimination at the midblastula transition in Drosophila melanogaster. EMBO J 18, 2610–2620.

Bawankar, P., Loh, B., Wohlbold, L., Schmidt, S., and Izaurralde, E. (2013). NOT10 and C2orf29/NOT11 form a conserved module of the CCR4-NOT complex that docks onto the NOT1 N-terminal domain. RNA Biol 10, 228–244.

Benoit, P., Papin, C., Kwak, J.E., Wickens, M., and Simonelig, M. (2008). PAP- and GLD-2-type poly(A) polymerases are required sequentially in cytoplasmic polyadenylation and oogenesis in Drosophila. Development 135, 1969–1979.

Berger, I., Fitzgerald, D.J., and Richmond, T.J. (2004). Baculovirus expression system for heterologous multiprotein complexes. Nat Biotechnol 22, 1583–1587.

Bhandari, D., Raisch, T., Weichenrieder, O., Jonas, S., and Izaurralde, E. (2014). Structural basis for the Nanos-mediated recruitment of the CCR4-NOT complex and translational repression. Genes Dev 28, 888–901.

Bhaskar, V., Roudko, V., Basquin, J., Sharma, K., Urlaub, H., Seraphin, B., and Conti, E. (2013). Structure and RNA-binding properties of the Not1-Not2Not5 module of the yeast Ccr4-Not complex. Nature Structural & Molecular Biology 20, 1281–1288.

Bienroth, S., Keller, W., and Wahle, E. (1993). Assembly of a Processive Messenger-Rna Polyadenylation Complex. Embo Journal 12, 585–594.

Boland, A., Chen, Y., Raisch, T., Jonas, S., Kuzuoglu-Ozturk, D., Wohlbold, L., Weichenrieder, O., and Izaurralde, E. (2013). Structure and assembly of the NOT module of the human CCR4-NOT complex. Nature Structural & Molecular Biology 20, 1289–1297.

Bosse-Doenecke, E., Weininger, U., Gopalswamy, M., Balbach, J., Knudsen, S.M., and Rudolph, R. (2008). High yield production of recombinant native and modified peptides exemplified by ligands for G-protein coupled receptors. Protein Expr Purif 58, 114–121.

Brandmann, T., Fakim, H., Padamsi, Z., Youn, J.Y., Gingras, A.C., Fabian, M.R., and Jinek, M. (2018). Molecular architecture of LSM14 interactions involved in the assembly of mRNA silencing complexes. Embo Journal 37, e97869.

Braun, J.E., Huntzinger, E., Fauser, M., and Izaurralde, E. (2011). GW182 proteins directly recruit cytoplasmic deadenylase complexes to miRNA targets. Mol Cell 44, 120–133.

Brawerman, G. (1981). The role of poly(A) sequence in mammalian messenger RNA. CRC Crit Rev Biochem 10, 1–38.

Brooks, S.A., and Blackshear, P.J. (2013). Tristetraprolin (TTP): Interactions with mRNA and proteins, and current thoughts on mechanisms of action. Bba-Gene Regul Mech 1829, 666–679.

Bulbrook, D., Brazier, H., Mahajan, P., Kliszczak, M., Fedorov, O., Marchese, F.P., Aubareda, A., Chalk, R., Picaud, S., Strain-Damerell, C., et al. (2018). Tryptophan-Mediated Interactions between Tristetraprolin and the CNOT9 Subunit Are Required for CCR4-NOT Deadenylase Complex Recruitment. J Mol Biol 430, 722–736.

Busso, D., Peleg, Y., Heidebrecht, T., Romier, C., Jacobovitch, Y., Dantes, A., Salim, L., Troesch, E., Schuetz, A., Heinemann, U., et al. (2011). Expression of protein complexes using multiple Escherichia coli protein co-expression systems: a benchmarking study. J Struct Biol 175, 159–170.

Cao, W.X., Kabelitz, S., Gupta, M., Yeung, E., Lin, S., Rammelt, C., Ihling, C., Pekovic, F., Low, T.C.H., Siddiqui, N.U., et al. (2020). Precise Temporal Regulation of Post-transcriptional Repressors Is Required for an Orderly Drosophila Maternal-to-Zygotic Transition. Cell Rep 31, 107783.

Chang, H., Lim, J., Ha, M., and Kim, V.N. (2014). TAIL-seq: Genome-wide Determination of Poly(A) Tail Length and 3 ‘ End Modifications. Molecular Cell 53, 1044–1052.

Charenton, C., and Graille, M. (2018). mRNA decapping: finding the right structures. Philos T R Soc B 373, 20180164.

Chartier, A., Klein, P., Pierson, S., Barbezier, N., Gidaro, T., Casas, F., Carberry, S., Dowling, P., Maynadier, L., Bellec, M., et al. (2015). Mitochondrial Dysfunction Reveals the Role of mRNA Poly(A) Tail Regulation in Oculopharyngeal Muscular Dystrophy Pathogenesis. PLoS Genet 11, e1005092.

Chekulaeva, M., Hentze, M.W., and Ephrussi, A. (2006). Bruno acts as a dual repressor of oskar translation, promoting mRNA oligomerization and formation of silencing particles. Cell 124, 521–533.

Chekulaeva, M., Mathys, H., Zipprich, J.T., Attig, J., Colic, M., Parker, R., and Filipowicz, W. (2011). miRNA repression involves GW182-mediated recruitment of CCR4-NOT through conserved W-containing motifs. Nat Struct Mol Biol 18, 1218–1226.

Chen, L., Dumelie, J.G., Li, X., Cheng, M.H., Yang, Z., Laver, J.D., Siddiqui, N.U., Westwood, J.T., Morris, Q., Lipshitz, H.D., et al. (2014a). Global regulation of mRNA translation and stability in the early Drosophila embryo by the Smaug RNA-binding protein. Genome Biol 15, R4.

Chen, Y., Boland, A., Kuzuoglu-Öztürk, D., Bawankar, P., Loh, B., Chang, C.-T., Weichenrieder, O., and Izaurralde, E. (2014b). A DDX6-CNOT1 complex and W-binding pockets in CNOT9 reveal direct links between miRNA target recognition and silencing. Mol Cell 54, 737–50.

Chen, Y., Khazina, E., Izaurralde, E., and Weichenrieder, O. (2021). Crystal structure and functional properties of the human CCR4-CAF1 deadenylase complex. Nucleic Acids Res 49, 6489–6510.

Colegrove-Otero, L.J., Minshall, N., and Standart, N. (2005). RNA-Binding proteins in early development. Crit Rev Biochem Mol Biol 40, 21–73.

Cooke, A., Prigge, A., and Wickens, M. (2010). Translational repression by deadenylases. J Biol Chem 285, 28506–28513.

Dahanukar, A., Walker, J.A., and Wharton, R.P. (1999). Smaug, a novel RNA-binding protein that operates a translational switch in Drosophila. Mol Cell 4, 209–218.

Dahanukar, A., and Wharton, R.P. (1996). The Nanos gradient in Drosophila embryos is generated by translational regulation. Genes Dev 10, 2610–2620.

Decker, C.J., and Parker, R. (1993). A Turnover Pathway for Both Stable and Unstable Messenger-Rnas in Yeast - Evidence for a Requirement for Deadenylation. Gene Dev 7, 1632–1643.

Diebold, M.L., Fribourg, S., Koch, M., Metzger, T., and Romier, C. (2011). Deciphering correct strategies for multiprotein complex assembly by co-expression: application to complexes as large as the histone octamer. J Struct Biol 175, 178–188.

Eckmann, C.R., Rammelt, C., and Wahle, E. (2011). Control of poly(A) tail length. Wires Rna 2, 348–361.

Eichhorn, S.W., Subtelny, A.O., Kronja, I., Kwasnieski, J.C., Orr-Weaver, T.L., and Bartel, D.P. (2016). mRNA poly(A)-tail changes specified by deadenylation broadly reshape translation in Drosophila oocytes and early embryos. Elife 5, e16955.

Eisen, T.J., Eichhorn, S.W., Subtelny, A.O., Lin, K.S., McGeary, S.E., Gupta, S., and Bartel, D. P. (2020). The Dynamics of Cytoplasmic mRNA Metabolism. Molecular Cell 77, 786–799.

Enwerem, I.I.I., Elrod, N.D., Chang, C.T., Lin, A., Ji, P., Bohn, J.A., Levdansky, Y., Wagner, E. J., Valkov, E., and Goldstrohm, A.C. (2021). Human Pumilio proteins directly bind the CCR4-NOT deadenylase complex to regulate the transcriptome. Rna 27, 445–464.

Ezzeddine, N., Chang, T.C., Zhu, W., Yamashita, A., Chen, C.Y., Zhong, Z., Yamashita, Y., Zheng, D., and Shyu, A.B. (2007). Human TOB, an antiproliferative transcription factor, is a poly(A)-binding protein-dependent positive regulator of cytoplasmic mRNA deadenylation. Mol Cell Biol 27, 7791–7801.

Fabian, M.R., Cieplak, M.K., Frank, F., Morita, M., Green, J., Srikumar, T., Nagar, B., Yamamoto, T., Raught, B., Duchaine, T.F., et al. (2011). miRNA-mediated deadenylation is orchestrated by GW182 through two conserved motifs that interact with CCR4-NOT. Nature Structural & Molecular Biology 18, 1211–1217.

Fabian, M.R., Frank, F., Rouya, C., Siddiqui, N., Lai, W.S., Karetnikov, A., Blackshear, P.J., Nagar, B., and Sonenberg, N. (2013). Structural basis for the recruitment of the human CCR4-NOT deadenylase complex by tristetraprolin. Nature Structural & Molecular Biology 20, 735–739.

Fitzgerald, D.J., Berger, P., Schaffitzel, C., Yamada, K., Richmond, T.J., and Berger, I. (2006). Protein complex expression by using multigene baculoviral vectors. Nat Methods 3, 1021–1032.

Garces, R.G., Gillon, W., and Pai, E.F. (2007). Atomic model of human Rcd-1 reveals an armadillo-like-repeat protein with in vitro nucleic acid binding properties. Protein Sci 16, 176–188.

Gavis, E.R., and Lehmann, R. (1992). Localization of nanos RNA controls embryonic polarity. Cell 71, 301–313.

Gavis, E.R., and Lehmann, R. (1994). Translational regulation of nanos by RNA localization. Nature 369, 315 – 318.

Gavis, E.R., Lunsford, L., Bergsten, S.E., and Lehmann, R. (1996). A conserved 90 nucleotide element mediates translational repression of nanos RNA. Development 122, 2791–2800.

Gerstberger, S., Hafner, M., and Tuschl, T. (2014). A census of human RNA-binding proteins. Nat Rev Genet 15, 829–845.

Goldstrohm, A.C., Hall, T.M.T., and McKenney, K.M. (2018). Post-transcriptional Regulatory Functions of Mammalian Pumilio Proteins. Trends in Genetics 34, 972–990.

Goldstrohm, A.C., and Wickens, M. (2008). Multifunctional deadenylase complexes diversify mRNA control. Nat Rev Mol Cell Bio 9, 337–344.

Götze, M., Dufourt, J., Ihling, C., Rammelt, C., Pierson, S., Sambrani, N., Temme, C., Sinz, A., Simonelig, M., and Wahle, E. (2017). Translational repression of the Drosophila nanos mRNA involves the RNA helicase Belle and RNA coating by Me31B and Trailer hitch. RNA 23, 1552–1568.

Green, J.B., Gardner, C.D., Wharton, R.P., and Aggarwal, A.K. (2003). RNA recognition via the SAM domain of Smaug. Mol Cell 11, 1537–1548.

Igreja, C., and Izaurralde, E. (2011). CUP promotes deadenylation and inhibits decapping of mRNA targets. Genes Dev 25, 1955–1967.

Iwasaki, S., Kawamata, T., and Tomari, Y. (2009). Drosophila argonaute1 and argonaute2 employ distinct mechanisms for translational repression. Mol Cell 34, 58–67.

Jeske, M., Bordi, M., Glatt, S., Muller, S., Rybin, V., Muller, C.W., and Ephrussi, A. (2015). The Crystal Structure of the Drosophila Germline Inducer Oskar Identifies Two Domains with Distinct Vasa Helicase- and RNA-Binding Activities. Cell Rep 12, 587–598.

Jeske, M., Meyer, S., Temme, C., Freudenreich, D., and Wahle, E. (2006). Rapid ATP-dependent deadenylation of nanos mRNA in a cell-free system from Drosophila embryos. J Biol Chem 281, 25124–25133.

Jeske, M., Moritz, B., Anders, A., and Wahle, E. (2011). Smaug assembles an ATP-dependent stable complex repressing nanos mRNA translation at multiple levels. EMBO J 30, 90–103.

Joly, W., Chartier, A., Rojas-Rios, P., Busseau, I., and Simonelig, M. (2013). The CCR4 deadenylase acts with Nanos and Pumilio in the fine-tuning of Mei-P26 expression to promote germline stem cell self-renewal. Stem Cell Reports 1, 411–424.

Kafasla, P., Mickleburgh, I., Llorian, M., Coelho, M., Gooding, C., Cherny, D., Joshi, A., Kotik-Kogan, O., Curry, S., Eperon, I.C., et al. (2012). Defining the roles and interactions of PTB. Biochem Soc T 40, 815–820.

Kamenska, A., Lu, W.T., Kubacka, D., Broomhead, H., Minshall, N., Bushell, M., and Standart, N. (2014). Human 4E-T represses translation of bound mRNAs and enhances microRNA-mediated silencing. Nucleic Acids Res 42, 3298–3313.

Kamenska, A., Simpson, C., Vindry, C., Broomhead, H., Benard, M., Ernoult-Lange, M., Lee, B.P., Harries, L.W., Weil, D., and Standart, N. (2016). The DDX6-4E-T interaction mediates translational repression and P-body assembly. Nucleic Acids Research 44, 6318–6334.

Kim-Ha, J., Kerr, K., and Macdonald, P.M. (1995). Translational Regulation of Oskar Messenger-Rna by Bruno, an Ovarian Rna-Binding Protein, Is Essential. Cell 81, 403–412.

Kinkelin, K., Veith, K., Grunwald, M., and Bono, F. (2012). Crystal structure of a minimal eIF4E-Cup complex reveals a general mechanism of eIF4E regulation in translational repression. RNA 18, 1624–1634.

Kluge, F., Gotze, M., and Wahle, E. (2020). Establishment of 5’-3’ interactions in mRNA independent of a continuous ribose-phosphate backbone. RNA 26, 613–628.

Kühn, U., Buschmann, J., and Wahle, E. (2017). The nuclear poly(A) binding protein of mammals, but not of fission yeast, participates in mRNA polyadenylation. Rna 23, 473–482.

Laver, J.D., Marsolais, A.J., Smibert, C.A., and Lipshitz, H.D. (2015). Regulation and Function of Maternal Gene Products During the Maternal-to-Zygotic Transition in Drosophila. Curr Top Dev Biol 113, 43–84.

Legnini, I., Alles, J., Karaiskos, N., Ayoub, S., and Rajewsky, N. (2019). FLAM-seq: fulllength mRNA sequencing reveals principles of poly(A) tail length control. Nat Methods 16, 879–886.

Leppek, K., Schott, J., Reitter, S., Poetz, F., Hammond, M.C., and Stoecklin, G. (2013). Roquin Promotes Constitutive mRNA Decay via a Conserved Class of Stem-Loop Recognition Motifs. Cell 153, 869–881.

Lim, J., Lee, M., Son, A., Chang, H., and Kim, V.N. (2016). mTAIL-seq reveals dynamic poly(A) tail regulation in oocyte-to-embryo development. Genes Dev 30, 1671–1682.

Lima, S.A., Chipman, L.B., Nicholson, A.L., Chen, Y.H., Yee, B.A., Yeo, G.W., Coller, J., and Pasquinelli, A.E. (2017). Short poly(A) tails are a conserved feature of highly expressed genes. Nature Structural & Molecular Biology 24, 1057–1063.

Maryati, M., Airhihen, B., and Winkler, G.S. (2015). The enzyme activities of Caf1 and Ccr4 are both required for deadenylation by the human Ccr4-Not nuclease module. Biochem J 469, 169–176.

Mathys, H., Basquin, J., Ozgur, S., Czarnocki-Cieciura, M., Bonneau, F., Aartse, A., Dziembowski, A., Nowotny, M., Conti, E., and Filipowicz, W. (2014). Structural and biochemical insights to the role of the CCR4-NOT complex and DDX6 ATPase in microRNA represson. Mol Cell 54, 751–765.

Mauxion, F., Prève, B., and Sèraphin, B. (2013). C2ORF29/CNOT11 and CNOT10 form a new module of the CCR4-NOT complex. RNA Biol 10, 267–276.

Mostafa, D., Takahashi, A., Yanagiya, A., Yamaguchi, T., Abe, T., Kureha, T., Kuba, K., Kanegae, Y., Furuta, Y., Yamamoto, T., et al. (2020). Essential functions of the CNOT7/8 catalytic subunits of the CCR4-NOT complex in mRNA regulation and cell viability. RNA Biol 17, 403–416.

Muhlrad, D., Decker, C.J., and Parker, R. (1994). Deadenylation of the unstable mRNA encoded by the yeast MFA2 gene leads to decapping followed by 5’-3’ digestion of the transcript. Genes Dev 8, 855–866.

Nakamura, A., Amikura, R., Hanyu, K., and Kobayashi, S. (2001). Me31B silences translation of oocyte-localizing RNAs through the formation of cytoplasmic RNP complex during Drosophila oogenesis. Development 128, 3233–3242.

Nakamura, A., Sato, K., and Hanyu-Nakamura, K. (2004). Drosophila cup is an eIF4E binding protein that associates with Bruno and regulates oskar mRNA translation in oogenesis. Dev Cell 6, 69–78.

Nelson, M.R., Leidal, A.M., and Smibert, C.A. (2004). Drosophila Cup is an eIF4E-binding protein that functions in Smaug-mediated translational repression. EMBO J 23, 150–159.

Niinuma, S., and Tomari, Y. (2017). ATP is dispensable for both miRNA- and Smaug-mediated deadenylation reactions. RNA 23, 866–871.

Nishimura, T., Padamsi, Z., Fakim, H., Milette, S., Dunham, W.H., Gingras, A.C., and Fabian, M. R. (2015). The eIF4E-Binding Protein 4E-T Is a Component of the mRNA Decay Machinery that Bridges the 5’ and 3’ Termini of Target mRNAs. Cell Rep 11, 1425–1436.

O’Farrell (2015). Growing an embryo from a single cell: A hurdle in animal life. Cold Spring Harb Perspect Biol 7, a019042.

Ozgur, S., Basquin, J., Kamenska, A., Filipowicz, W., Standart, N., and Conti, E. (2015). Structure of a Human 4E-T/DDX6/CNOT1 Complex Reveals the Different Interplay of DDX6-Binding Proteins with the CCR4-NOT Complex. Cell Rep 13, 703–711.

Park, E.H., Walker, S.E., Lee, J.M., Rothenburg, S., Lorsch, J.R., and Hinnebusch, A.G. (2011). Multiple elements in the eIF4G1 N-terminus promote assembly of elF4G1 center dot PABP mRNPs in vivo. Embo Journal 30, 302–316.

Pavanello, L., Hall, B., Airhihen, B., and Winkler, G.S. (2018). The central region of CNOT1 and CNOT9 stimulates deadenylation by the Ccr4-Not nuclease module. Biochem J 475, 3437–3450.

Raisch, T., Bhandari, D., Sabath, K., Helms, S., Valkov, E., Weichenrieder, O., and Izaurralde, E. (2016). Distinct modes of recruitment of the CCR4-NOT complex by Drosophila and vertebrate Nanos. EMBO J 35, 974–990.

Raisch, T., Chang, C.T., Levdansky, Y., Muthukumar, S., Raunser, S., and Valkov, E. (2019). Reconstitution of recombinant human CCR4-NOT reveals molecular insights into regulated deadenylation. Nat Commun 10, 3173.

Räsch, F., Weber, R., and Igreja, C. (2020). 4E-T-bound mRNAs are stored in a silenced and deadenylated form. Gene Dev 34, 847–860.

Richter, J.D. (2000). Influence of polyadenylation-induced translation on metazoan development and neuronal synaptic function. In Translational control of gene expression, N. Sonenberg, J.W.B. Hershey, and M.B. Mathews, eds. (Cold Spring Harbor, New York: Cold Spring Harbor Laboratory Press).

Rouget, C., Papin, C., Boureux, A., Meunier, A.C., Franco, B., Robine, N., Lai, E.C., Pelisson, A., and Simonelig, M. (2010). Maternal mRNA deadenylation and decay by the piRNA pathway in the early Drosophila embryo. Nature 467, 1128–1132.

Rouya, C., Siddiqui, N., Morita, M., Duchaine, T.F., Fabian, M.R., and Sonenberg, N. (2014). Human DDX6 effects miRNA-mediated gene silencing via direct binding to CNOT1. RNA 20, 1398–1409.

Salgania, H.K., Metz, J., and Jeske, M. (2022). ReLo: a simple colocalization assay to identify and characterize physical protein-protein interactions. BioRxiv, doi: 10.1101/2022.1103.1104.482790

Salles, F.J., Lieberfarb, M.E., Wreden, C., Gergen, J.P., and Strickland, S. (1994). Coordinate Initiation of Drosophila Development by Regulated Polyadenylation of Maternal Messenger-Rnas. Science 266, 1996–1999.

Sawicki, S.G., Jelinek, W., and Darnell, J.E. (1977). 3’-Terminal Addition to Hela-Cell Nuclear and Cytoplasmic Poly(a). J Mol Biol 113, 219–235.

Schindelin, J., Arganda-Carreras, I., Frise, E., Kaynig, V., Longair, M., Pietzsch, T., Preibisch, S., Rueden, C., Saalfeld, S., Schmid, B., et al. (2012). Fiji: an open-source platform for biological-image analysis. Nat Methods 9, 676–682.

Semotok, J.L., Cooperstock, R.L., Pinder, B.D., Vari, H.K., Lipshitz, H.D., and Smibert, C.A. (2005). Smaug recruits the CCR4/POP2/NOT deadenylase complex to trigger maternal transcript localization in the early Drosophila embryo. Curr Biol 15, 284–294.

Semotok, J.L., and Lipshitz, H.D. (2007). Regulation and function of maternal mRNA destabilization during early Drosophila development. Differentiation 75, 482–506.

Sgromo, A., Raisch, T., Bawankar, P., Bhandari, D., Chen, Y., Kuzuoglu-Ozturk, D., Weichenrieder, O., and Izaurralde, E. (2017). A CAF40-binding motif facilitates recruitment of the CCR4-NOT complex to mRNAs targeted by Drosophila Roquin. Nat Commun 8, 14307.

Sheu-Gruttadauria, J., and MacRae, I.J. (2018). Phase Transitions in the Assembly and Function of Human miRISC. Cell 173, 946–957.

Smibert, C.A., Lie, Y.S., Shillinglaw, W., Henzel, W.J., and Macdonald, P.M. (1999). Smaug, a novel and conserved protein, contributes to repression of nanos mRNA translation in vitro. RNA 5, 1535–1547.

Smibert, C.A., Wilson, J.E., Kerr, K., and Macdonald, P.M. (1996). Smaug protein represses translation of unlocalized nanos mRNA in the Drosophila embryo. Genes Dev 10, 2600–2609.

Stagljar, I., Korostensky, C., Johnsson, N., and te Heesen, S. (1998). A genetic system based on split-ubiquitin for the analysis of interactions between membrane proteins in vivo. Proc Natl Acad Sci U S A 95, 5187–5192.

Stowell, J.A.W., Webster, M.W., Kogel, A., Wolf, J., Shelley, K.L., and Passmore, L.A. (2016). Reconstitution of Targeted Deadenylation by the Ccr4-Not Complex and the YTH Domain Protein Mmi1. Cell Reports 17, 1978–1989.

Studier, F.W. (2005). Protein production by auto-induction in high density shaking cultures. Protein Expr Purif 41, 207–234.

Stukenberg, P.T., Studwell-Vaughan, P.S., and Odonnell, M. (1991). Mechanism of the Sliding Beta-Clamp of DNA Polymerase-Iii Holoenzyme. J. Biol. Chem. 266, 11328–11334.

Stupfler, B., Birck, C., Seraphin, B., and Mauxion, F. (2016). BTG2 bridges PABPC1 RNA-binding domains and CAF1 deadenylase to control cell proliferation. Nat Commun 7, 10811.

Subtelny, A.O., Eichhorn, S.W., Chen, G.R., Sive, H., and Bartel, D.P. (2014). Poly(A)-tail profiling reveals an embryonic switch in translational control. Nature 508, 66–71.

Sysoev, V.O., Fischer, B., Frese, C.K., Gupta, I., Krijgsveld, J., Hentze, M.W., Castello, A., and Ephrussi, A. (2016). Global changes of the RNA-bound proteome during the maternal-to-zygotic transition in Drosophila. Nat Commun 7, 12128.

Tadros, W., Goldman, A.L., Babak, T., Menzies, F., Vardy, L., Orr-Weaver, T., Hughes, T.R., Westwood, J.T., Smilbert, C.A., and Lipshitz, H.D. (2007). SMAUG is a major regulator of maternal mRNA destabilization in Drosophila and its translation is activated by the PAN GU kinase. Dev Cell 12, 143–155.

Tarun, S.Z., and Sachs, A.B. (1996). Association of the yeast poly(A) tail binding protein with translation initiation factor eIF-4G. EMBO J 15, 7168–7177.

Temme, C., Simonelig, M., and Wahle, E. (2014). Deadenylation of mRNA by the CCR4-NOT complex in Drosophila: molecular and developmental aspects. Front Genet 5, doi: 10.3389/fgene.2014.00143

Temme, C., Zhang, L.B., Kremmer, E., Ihling, C., Chartier, A., Sinz, A., Simonelig, M., and Wahle, E. (2010). Subunits of the Drosophila CCR4-NOT complex and their roles in mRNA deadenylation. Rna 16, 1356–1370.

Tritschler, F., Braun, J.E., Eulalio, A., Truffault, V., Izaurralde, E., and Weichenrieder, O. (2009). Structural basis for the mutually exclusive anchoring of P body components EDC3 and Tral to the DEAD box protein DDX6/Me31B. Mol Cell 33, 661–668.

Tritschler, F., Eulalio, A., Helms, S., Schmidt, S., Coles, M., Weichenrieder, O., Izaurralde, E., and Truffault, V. (2008). Similar modes of interaction enable Trailer Hitch and EDC3 to associate with DCP1 and Me31B in distinct protein complexes. Mol Cell Biol 28, 6695–6708.

Tucker, M., Valencia-Sanchez, M.A., Staples, R.R., Chen, J., Denis, C.L., and Parker, R. (2001). The transcription factor associated Ccr4 and Caf1 proteins are components of the major cytoplasmic mRNA deadenylase in Saccharomyces cerevisiae. Cell 104, 377–386.

Van Etten, J., Schagat, T.L., Hrit, J., Weidmann, C.A., Brumbaugh, J., Coon, J.J., and Goldstrohm, A.C. (2012). Human Pumilio proteins recruit multiple deadenylases to efficiently repress messenger RNAs. J Biol Chem 287, 36370–36383.

Waghray, S., Williams, C., Coon, J.J., and Wickens, M. (2015). Xenopus CAF1 requires NOT1-mediated interaction with 4E-T to repress translation in vivo. RNA 21, 1335–1345.

Wahle, E., and Winkler, G.S. (2013). RNA decay machines: Deadenylation by the Ccr4-Not and Pan2-Pan3 complexes. Biochim Biophys Acta 1829, 561–570.

Walser, C.B., and Lipshitz, H.D. (2011). Transcript clearance during the maternal-to-zygotic transition. Curr Opin Genet Dev 21, 431–443.

Wang, C., and Lehmann, R. (1991). Nanos is the localized posterior determinant in Drosophila. Cell 66, 637–647.

Webster, M.W., Chen, Y.H., Stowell, J.A.W., Alhusaini, N., Sweet, T., Graveley, B.R., Coller, J., and Passmore, L.A. (2018). mRNA Deadenylation Is Coupled to Translation Rates by the Differential Activities of Ccr4-Not Nucleases. Molecular Cell 70, 1089–1100.

Webster, M.W., Stowell, J.A.W., and Passmore, L.A. (2019). RNA-binding proteins distinguish between similar sequence motifs to promote targeted deadenylation by Ccr4-Not. Elife 8, e40670.

Wessels, H.H., Imami, K., Baltz, A.G., Kolinski, M., Beldovskaya, A., Selbach, M., Small, S., Ohler, U., and Landthaler, M. (2016). The mRNA-bound proteome of the early fly embryo. Genome Res 26, 1000–1009.

Wickens, M., Bernstein, D.S., Kimble, J., and Parker, R. (2002). A PUF family portrait: 3 ‘ UTR regulation as a way of life. Trends in Genetics 18, 150–157.

Wilhelm, J.E., Hilton, M., Amos, Q., and Henzel, W.J. (2003). Cup is an elF4E binding protein required for both the translational repression of oskar and the recruitment of Barentsz. Journal of Cell Biology 163, 1197–1204.

Wilson, T., and Treisman, R. (1988). Removal of Poly(a) and Consequent Degradation of CFos Messenger-Rna Facilitated by 3’ Au-Rich Sequences. Nature 336, 396–399.

Wu, L.G., Fan, J.H., and Belasco, J.G. (2006). MicroRNAs direct rapid deadenylation of mRNA. P Natl Acad Sci USA 103, 4034–4039.

Xiang, K., and Bartel, D.P. (2021). The molecular basis of coupling between poly(A)-tail length and translational efficiency. Elife 10, e66493.

Yi, H., Park, J., Ha, M., Lim, J., Chang, H., and Kim, V.N. (2018). PABP Cooperates with the CCR4-NOT Complex to Promote mRNA Deadenylation and Block Precocious Decay. Molecular Cell 70, 1081–1088.

Zaessinger, S., Busseau, I., and Simonelig, M. (2006). Oskar allows nanos mRNA translation in Drosophila embryos by preventing its deadenylation by Smaug/CCR4. Development 133, 4573–4583.

Zappavigna, V., Piccioni, F., Villaescusa, J.C., and Verrotti, A.C. (2004). Cup is a nucleocytoplasmic shuttling protein that interacts with the eukaryotic translation initiation factor 4E to modulate Drosophila ovary development. P Natl Acad Sci USA 101, 14800–14805.

