## Supplemental Figures and Tables for "RNA binding proteins Smaug and Cup induce CCR4-NOT-dependent deadenylation of the *nanos* mRNA in a reconstituted system"

Inventory:

Supplemental Figure 1
Supplemental Figure 2
Supplemental Figure 3
Supplemental Figure 4
Supplemental Figure 5
Supplemental Table 1
Supplemental Table 2

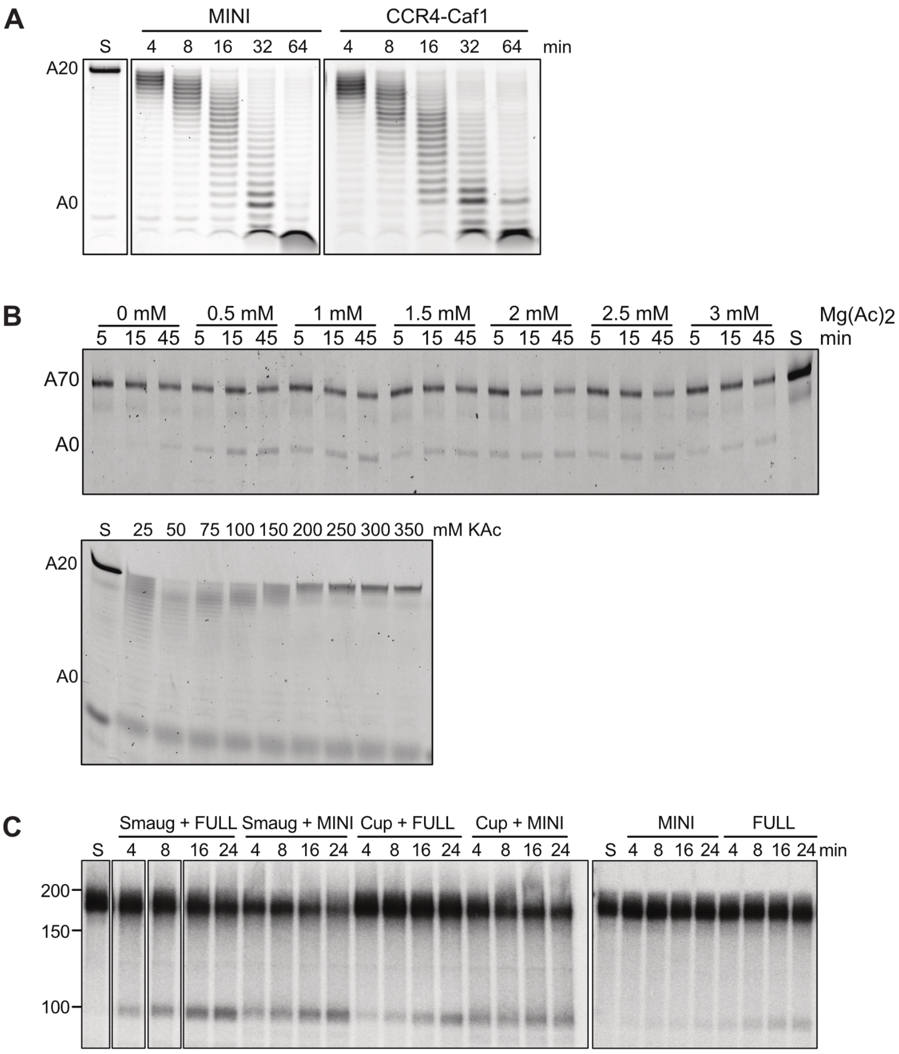


**Supplemental Figure 1. Activity of CCR4 and CAF1 depends on other subunits of CCR4-NOT and on ionic conditions**

(A) Deadenylation activity of the CCR4-Caf1 heterodimer is enhanced by other subunits of the CCR4-NOT complex. 50 nM FAM-7mer-A$_{20}$ RNA was incubated with either 5 nM of $^{Dm}$CCR4-NOT$_{MINI}$ or 250 nM of $^{Dm}$CCR4-Caf1, and aliquots were withdrawn as indicated. At the 50fold higher concentration, CCR4-Caf1 shows activity comparable to that of the CCR4-NOT$_{MINI}$ complex.

(B) Deadenylation activity of $^{Dm}$CCR4-NOT complex is sensitive to Mg$^{2+}$ and K$^{+}$ concentrations. Top panel: 25 nM FAM-TCE$^{MUT}$-A$_{70}$ RNA was incubated with 25 nM of $^{Dm}$CCR4-NOT$_{MINI}$ in the presence of varying concentrations of magnesium acetate. Aliquots were withdrawn and analyzed at the time points indicated. In this assay, relatively high concentrations of enzyme were used as stability of the CCR4-NOT complex had not yet been optimized. Bottom panel: 10 nM FAM-7mer-A$_{20}$ RNA was incubated with 5 nM of $^{Dm}$CCR4-NOT$_{MINI}$ in the presence of increasing concentrations of potassium acetate. These assays were done in the absence of BSA. Deadenylation was stopped after 60 minutes. In addition to the varying concentrations of potassium acetate, the reactions contained 3 mM NaCl introduced with the protein buffer. The strong diffuse band visible at the bottom is bromophenol blue.

(C) $^{Dm}$CCR4-NOT$_{FULL}$ and $^{Dm}$CCR4-NOT$_{MINI}$ behave similarly in Smaug- or Cup-dependent deadenylation. Reactions contained 20 nM SRE$^{WT}$-A$_{70}$ RNA, 1 nM of the CCR4-NOT complex indicated and 80 nM Smaug or Cup where indicated. Aliquots were withdrawn and analyzed at the time points indicated. S, unreacted substrate.

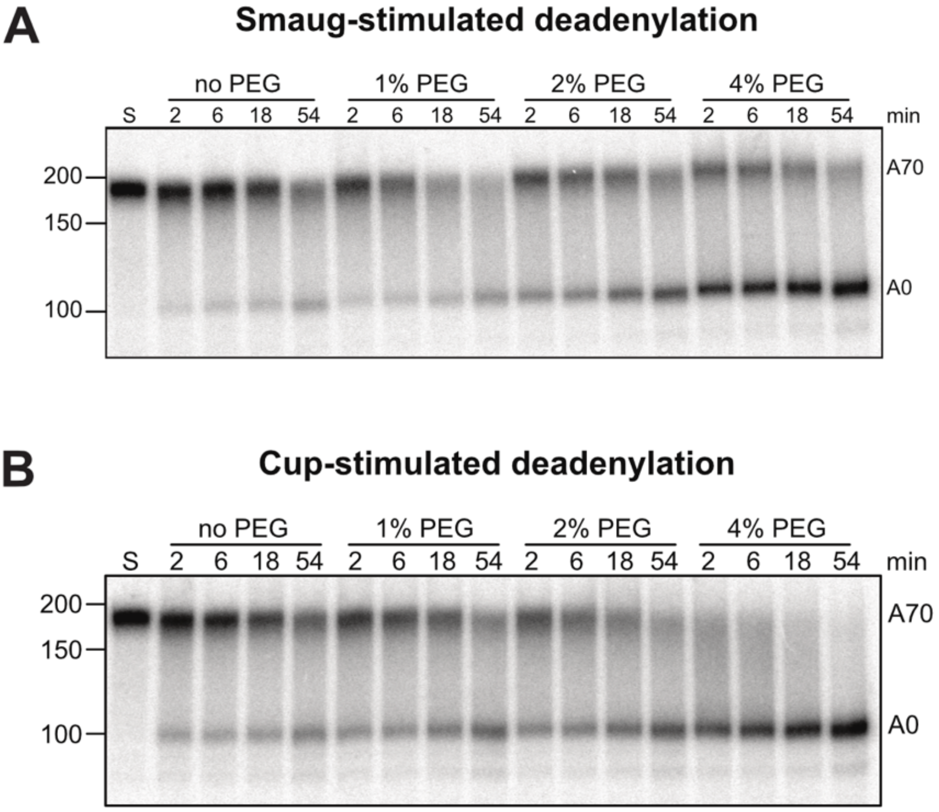


**Supplemental Figure 2. Smaug- and Cup-dependent deadenylation is enhanced by a crowding reagent**

(A) Smaug-dependent deadenylation is stimulated by a crowding reagent. Deadenylation time courses were carried out with 10 nM $SRE^{WT}$-$A_{70}$ RNA, 80 nM Smaug and 2 nM $^{Dm}CCR4\text{-}NOT_{MINI}$ in the presence of different concentrations of PEG 20,000.

(B) Cup-dependent deadenylation is stimulated by a crowding reagent. Deadenylation time courses were carried out as in (A) except that 80 nM Cup replaced Smaug.

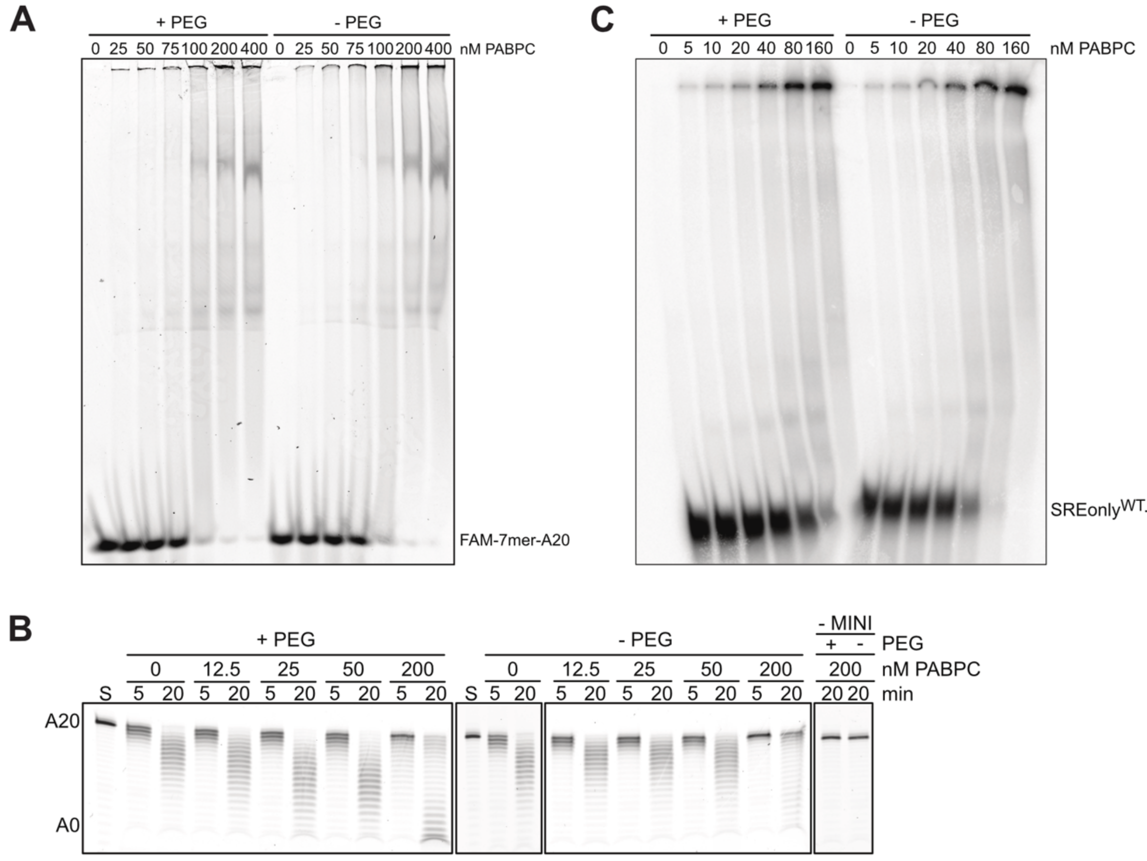


**Supplemental Figure 3. PABPC modestly stimulates deadenylation**

(A) Binding of PABPC to the FAM-7mer-$A_{20}$ RNA. Indicated concentrations of PABPC were incubated with 25 nM FAM 7mer-$A_{20}$ RNA at 25 °C for 20 minutes in the presence or absence of PEG. Then, RNA-protein complexes were separated by electrophoresis on a nondenaturing polyacrylamide gel run at 200 V and room temperature.

(B) PABPC stimulates deadenylation of FAM 7mer. FAM 7mer-$A_{20}$ RNA (25 nM) was incubated with the indicated concentrations of PABPC in the presence or absence of PEG. In these assays, tRNA was present as in the binding assay (**panel A**). The deadenylation reaction was started with $^{Dm}$CCR4-NOT$_{MINI}$ (1 nM) and was allowed to proceed for 5 or 20 min as indicated.

(C) Binding of PABPC to SRE$^{WT}$only-$A_{70}$ RNA. SRE$^{WT}$only-$A_{70}$ RNA (5 nM) were incubated with the indicated concentrations of PABPC in the presence or absence of PEG, and RNA-protein complexes were analyzed as in **(A)**. With an $A_{70}$ tail and one molecule of PABPC covering ~27 nt, saturation of the substrate RNA would have been expected at ~15 nM PABPC.

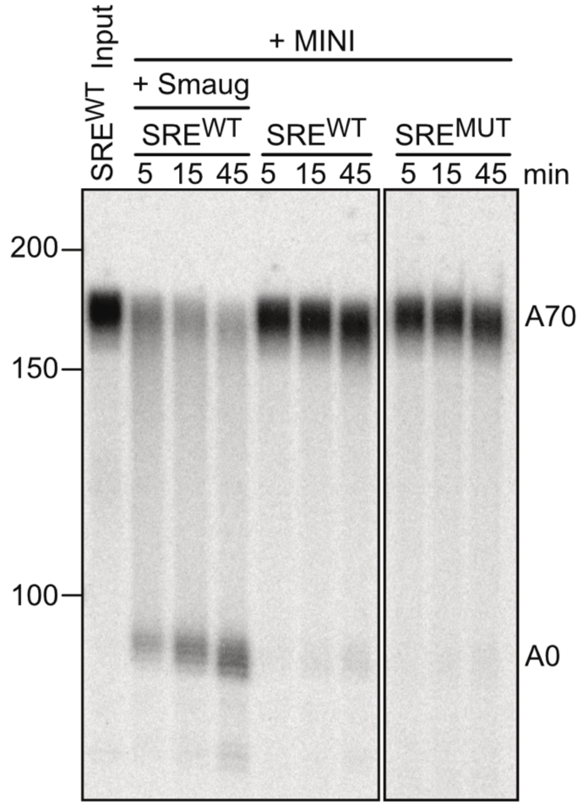


**Supplemental Figure 4. Smaug-independent deadenylation of a long substrate RNA is distributive**

20 nM SRE$^{WT}$only-$A_{70}$ RNA was preincubated for 20 min with 80 nM Smaug or buffer. SRE$^{MUT}$only-$A_{70}$ RNA was preincubated with buffer. Deadenylation was then initiated by the addition of 5 nM $^{Dm}$CCR4-NOT$_{MINI,}$ and aliquots were withdrawn at the time points indicated. Co-existence of full-length RNA and completely deadenylated product in the Smaug-containing reaction confirms processive activity, as in **Fig. 6B**. In the absence of Smaug, modest shortening of the entire population of substrate RNA, mostly visible at the latest time point, indicates weak, distributive activity. In this reaction, very small amounts of completely deadenylated product are also visible. These are no longer present when an SRE$^{MUT}$ substrate is used, suggesting the possibility that the CCR4-NOT preparation is contaminated by small amounts of a Smaug-like protein.

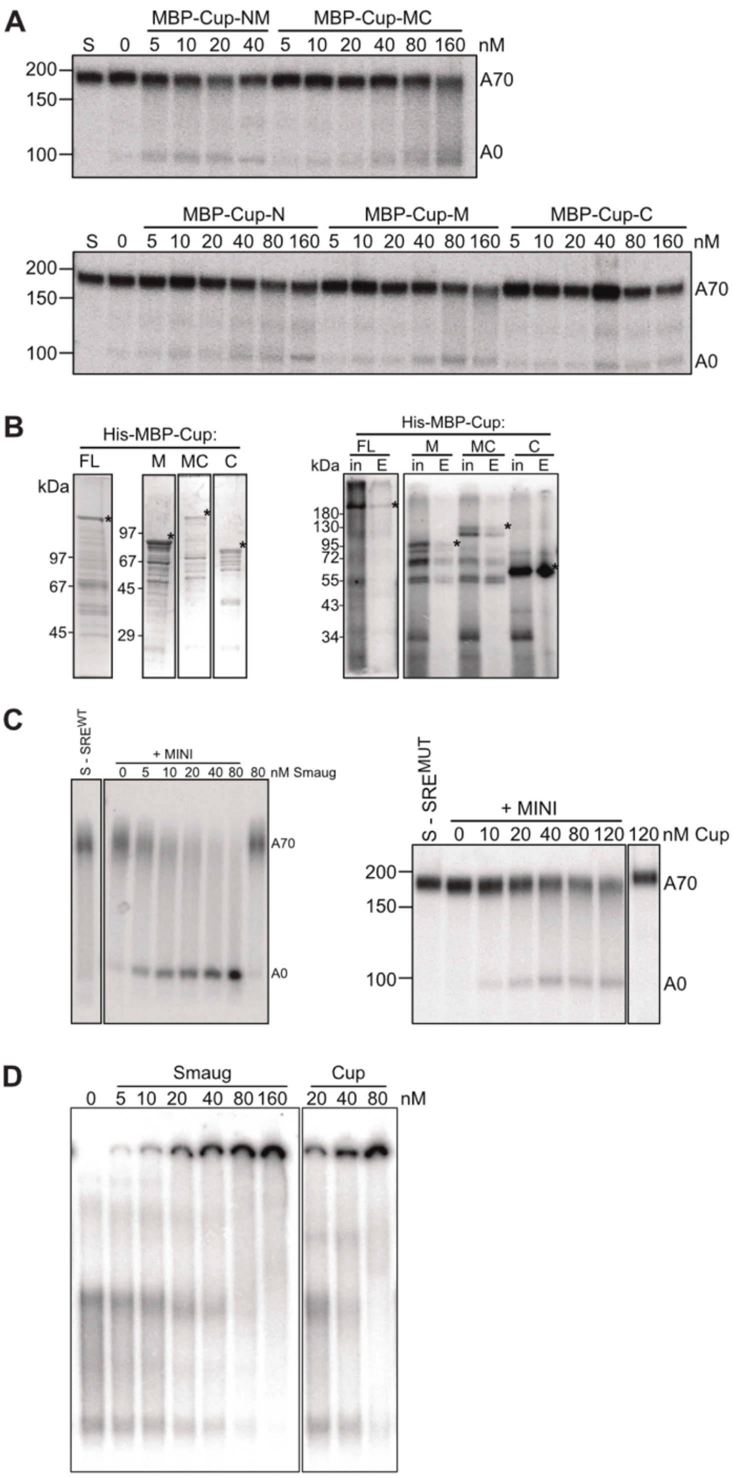


**Supplemental Figure 5. Cup contributes to deadenylation**

(A) The ability of Cup to stimulate deadenylation is distributed over the protein. 5 nM $^{32}$P-SRE$^{MUT}$only-A$_{70}$ RNA was preincubated in the presence of varying concentrations of different Cup fragments or in their absence, as indicated. Deadenylation was started by the addition of 1 nM $^{Dm}$CCR4-NOT$_{MINI}$. Reactions were allowed to proceed for 30 min. Controls in a separate experiment showed that the Cup fragments were devoid of nuclease activity, i. e. deadenylation was CNOT-dependent.

**Supplemental Figure 5 (continued). Cup contributes to deadenylation**

(B) Fragments of Cup can be UV cross-linked to RNA. Left panel, Coomassie-stained SDS-polyacrylamide gel lanes showing His-MBP-tagged Cup and Cup fragments. Bands corresponding to the desired proteins are marked. Right panel: Proteins shown in the left panel were UV-cross-linked to RNA (in, input fraction; 30 % of total reaction). 70% of each reaction were incubated with Ni-NTA matrix in the presence of 6M urea and 0.05 % NP-40, and bound protein-RNA complexes were eluted (E, eluate). Input and eluate were analyzed by SDS-polyacrylamide gel electrophoresis and autoradiography. Bands corresponding to the proteins of interest are labeled. (C) Titration of Smaug (left panel) and Cup (right panel) in deadenylation. 20 nM SREonly-$A_{70}$ RNA, wild-type or mutant as indicated, was preincubated for 30 min with the indicated concentrations of Cup or Smaug, respectively, then deadenylation was started by the addition of 2 nM $^{Dm}$CCR4-NOT$_{MINI}$ and stopped after 40 min. PEG was present in the Smaug titration, but not in the Cup titration.

(D) Titration of Smaug and Cup in RNA binding. 10 nM SRE$^{WT}$only-A70 RNA was incubated at 25 °C for 20 minutes with the indicated concentrations of Smaug or Cup in the absence of PEG. Then, RNA-protein complexes were separated by electrophoresis on a non-denaturing polyacrylamide gel. Retarded RNA-protein complexes were stuck in the wells.

**Supplemental Table 1. Sequences of synthetic oligonucleotides**
Capital letters indicate the start of the corresponding gene. Point mutations are indicated with bold letters.

| Gene etc. | Forward sequence (5'-3') | Reverse sequence (5'-3') | Plasmid |
|---|---|---|---|
| NOT1 PC | gctctagaATGGCTAGTAACGTAGAGAGCCAACTG | TGGCTAATAGCACGCGCTATC | pFBDM-His-NOT2_NOT1(Full) |
| NOT1 PE | gctctagaATGGCTAGTGACACATCTTGGATTAATC | TGGCTAATAGCACGCGCTATC | pFBDM-His-NOT2_NOT1(PE) |
| NOT1 MINI | ggactagtatgcatcatcaccatcaccatcaccatGTGACTGTGCCACCAGAG | cccaagcttTCAGTTGATGGTGGCTAC | pET28a-MBP-His8-NOT1(MINI) |
| | | | pFBDM-His8-NOT2_His8-NOT1(MINI) |
| NOT2 | tcccccgggatgcatcatcaccatcaccatcaccatATGGCGAATTTAAATTTTC | ggggtaccTTATACAGACTGTCCATTC | pFBDM-His8-NOT2_His8-NOT1(MINI) |
| NOT3 | tcccccgggatgcatcatcaccatcaccatcaccatATGGCTGCGACGAGAAAATTG | ggggtaccTCAATTCAGCTCCTTGTC | pFBDM-CAF40-FLAG_His8-NOT3 |
| Caf40 | ggaattcATGAGTGCTCAACCGAGTC | cccaagcttctacttatcgtcgtcatccttgtaatcGGAGCCCAGTGGCGACATG | pFBDM-CAF40-FLAG_His8-NOT3 |
| CAF1 | gcgcggatccatgtctcatcatcatcatcatcatcaccacATCAAATGGACAATGCCC | gctctagaTCATGAAGCGCTGTTCGTC | pFBDM-CCR4-FLAG_His8-CAF1 |
| CCR4 | ggatctcgagATGAAAGGCAATCATTATAAA | tcccggtaccttaatccttgtcatcgtcgtccttgtagtcGGCCCGGCGATTGATCAGC | pFBDM-CCR4-FLAG_His8-CAF1 |
| CAF1 (MUT) | CACTATGTGGCCATGG**c**CACCG**c**GTTTCCAGGCGTGGTAG | CTACCACGCCTGGAAAC**g**CGGTG**g**CCATGGCCACATAGTG | pFBDM-CCR4-FLAG_His8-CAF1(MUT) |
| | | | pFBDM-CCR4-FLAG(MUT)_His8-CAF1(MUT) |
| CCR4 (MUT) | GCTGCTGCTGTGCGGTG**c**CTTC**gc**CTCGCTACCCGATTCAG | CTGAATCGGGTAGCGAG**gc**GAAG**g**CACCGCACAGCAGCAGC | pFBDM-CCR4-FLAG(MUT)_His8-CAF1 |
| | | | pFBDM-CCR4-FLAG(MUT)_His8-CAF1(MUT) |
| Smaug | ctagctagcATGAAGTACGCAACTGGAAC | ataagaatgcggccgcctatttatcatcatcatctttataatcGAATAGCGTAAAATGTTG | pFBDM-Smaug-FLAG |
| Cup | tcccccgggctttattttcagggc | ccgctcgagttacttatcgtcgtcatccttgtaatcATGAAACTCATCCCCGC | pFBDM-Cup-FLAG |
| Tral | cggaattcATGAGCGGGGGATTACCG | atagccatggtcagtggtgatgatgatgatgatgatgTTGTGAAACTGCCGCCAC | pFBDM-Tral-His8 |

| | | | |
|---|---|---|---|
| PABPC | cgcggatccatgcatcatcatcatcatcatcaccacATGGCTTCTCTATACGTC | ccgctcgagTTAGTTGGCGGGCTCGGTG | pET28a-PABPC |
| Belle | tgcagtctcgagatgcatcatcatcatcatcatcaccacAGTAATGCTATTAACC | gtcgacaagcttTCATTGAGCCCACCA | pFastBac1-His8-Belle |
| CupNM | ccgctcgagATGCAAATGGCCGAAGC | cgcctaggTTATCGACGCCATTTG | pnEK-His8-MBP-CupNM |
| CupMC | ccgctcgagGACGAGTCCATC | ccgcctaggTTAATGAAACTCATCC | pnEK-His8-MBP-CupMC |
| CupN | ccgctcgagATGCAAATGGCCGAAGC | cgcctaggTTAGTCACTGATTAGGTTC | pnEK-His8-MBP-CupN |
| CupM | ccgctcgagGACGAGTCCATC | cgcctaggTTATCGACGCCATTTG | pnEK-His8-MBP-CupM |
| CupC | ccgctcgagCGAAACTCACTGAAC | ccgcctaggTTAATGAAACTCATCC | pnEK-His8-MBP-CupC |
| SmaugNM | cgggatccATGAAGTACGCAACTGGAAC | tcccccgggTTAAATATTGGCCCGTTCCTTC | pET28a-SmaugNM |
| SmaugMC | cgggatccTGCCCCGCAAGCGGCAG | tcccccgggTTAGAATAGCGTAAAATGTTG | pGEX6p1-SmaugMC |
| SmaugN | cgggatccATGAAGTACGCAACTGGAAC | tcccccgggTTACAACGAGGATGAGGCCAC | pGEX6p1-SmaugN |
| SmaugM | atatggtctcatggtAATTATATTAAGTTCCACACGCGC | tatctcgagttaATTATTCAGCGACCGGC | pET-SUMOadapt-SmaugM |
| SmaugC | cggaattcCTTAACCGGGTAGAACAAG | tcccccgggTTAGAATAGCGTAAAATGTTG | pGEX6p1-SmaugC |
| His-tag | catgggcagcagccatcatcaccatcaccatcaccattc | catggaatggtgatggtgatggtgatgatggctgctgcc | pnEK-His8-MBP |
| 2xBoxB | gatccGGGCCCTGAAGAAGGGCCCATATAGGGCCCTGAAGAAGGGCCCt (BamHI fragment) | ctagaGGGCCCTTCTTCAGGGCCCTATATGGGCCCTTCTTCAGGGCCCg (XbaI fragment) | pBSK-nLuc-2xBoxB |
| $BRE^{WT}$ | ggaagatctGAATTCGCTTAGTTTTAATATG | gctctagaggatccTTAAATCTAACATAGAAC | pBSK-nLuc-$BRE^{WT}$ |
| $BRE^{MUT}$ EcoRI | AATTCGCTTAGTTTTAAT**ta**GTTTT**ta**AT**tg**AG**at**TGTTCTCTGTCTTTGTT**at**TTTA**g**AT**t**TTCGTGCACTT (EcoRI fragment1) | AA**a**AT**c**TAAA**at**AACAAAGACAGAGAACA**at**CT**ca**AT**ta**AAAAC**ta**ATTAAAACTAAGCG (EcoRI fragment2) | pBSK-nLuc-$BRE^{MUT}$ |
| $BRE^{MUT}$BamHI | GTCCTAGTCCATTATT**t**TA**g**ATTATT**t**TG**g**GTTTTG**gt**TTCT**ta**GTTAGATTTAAG (BamHI fragment1) | GATCCTTAAATCTAAC**ta**AGAA**ac**CAAAAC**c**CA**a**AATAAT**c**TA**a**AATAATGGACTAGGACAAGTGCACG (BamHI fragment2) | |

**Supplemental Table 2. ReLo and Y2H DNA constructs used in this study.**
All bait and prey sequences were from *Drosophila melanogaster*.
The split-ubiquitin Y2H cloning vectors pDHB1-MJ (JK16) and pPR3-N-MJ (JK18) were generated by introducing the blunt end restriction sites Eco47III and SmaI into the multiple cloning sites of pDHB1 and pPR3-N (Jeske et al., 2015), respectively. The plasmids pAc5.1-EGFP (T5-MJ), pAc5.1-mCherry (T7-MJ), pAc5.1-PH-mCherry-CAF1 (HK96), pAc5.1-PH-mCherry-CAF40 (HK97), pAc5.1-PH-mCherry-NOT2 (HK99), pAc5.1-PH-mCherry-NOT3 (HK100), pAc5.1-CCR4-mCherry-PH (EB7), and pAc5.1-NOT1-mCherry-PH (EB5) have been described previously (Salgania et al., 2022). Generation of the DNA constructs listed in the table was performed according to the cloning strategy described previously (Salgania et al., 2022).

| **Vector**<br>(insertion site) (code) | **Final DNA construct** | **DNA template information** | **Code** |
|---|---|---|---|
| **pAc5.1-EGFP**<br>(EcoRV) (T5-MJ) | pAc5.1-EGFP-**Smaug** | *smaug* cDNA | F31-MJ |
| **pAc5.1-mCherry**<br>(EcoRV) (T7-MJ) | pAc5.1-mCherry-**Smaug** | *smaug* cDNA | H28-MJ |
| **pAc5.1-PH-mEGFP**<br>(FspAI) (JM50) | pAc5.1-PH-mEGFP-**NOT3 1-241** | pFL-Flag-NOT3 | JM69 |
| | pAc5.1-PH-mEGFP-**NOT3 242-686** | pFL-Flag-NOT3 | JM70 |
| | pAc5.1-PH-mEGFP-**NOT3 687-844** | pFL-Flag-NOT3 | JM71 |
| **pDHB1-MJ**<br>(Eco47III) (JK16) | pDHB1-MJ-**Smaug** | *smaug* cDNA | JK46 |
| **pPR3-N-MJ**<br>(SmaI) (JK18) | pPR3-N-MJ-**CAF1** | pMTV5-Myc-CAF1 (Temme et al., 2010) | JK76 |
| | pPR3-N-MJ-**CAF40** | pET19-CAF40 | JK77 |
| | pPR3-N-MJ-**CCR4** | pMTV5-Myc-CCR4 | JK78 |
| | pPR3-N-MJ-**NOT2** | pSPL_Strep_NOT1_NOT2 | JK88 |
| | pPR3-N-MJ-**NOT3** | pFL-Flag-NOT3 | JK89 |
| | pPR3-N-MJ-**NOT1 1-751** | pSPL_Strep_NOT1_NOT2 | JK84 |
| | pPR3-N-MJ-**NOT1 752-910** | pSPL_Strep_NOT1_NOT2 | FH16 |
| | pPR3-N-MJ-**NOT1 911-1092** | pSPL_Strep_NOT1_NOT2 | FH13 |
| | pPR3-N-MJ-**NOT1 1093-1687** | pSPL_Strep_NOT1_NOT2 | JK86 |
| | pPR3-N-MJ-**NOT1 1688-1964** | pSPL_Strep_NOT1_NOT2 | FH4 |
| | pPR3-N-MJ-**NOT1 1965-2480** | pSPL_Strep_NOT1_NOT2 | JK87 |
| | pPR3-N-MJ-**NOT3 1-241** | pFL-Flag-NOT3 | MS5 |
| | pPR3-N-MJ-**NOT3 242-686** | pFL-Flag-NOT3 | MS9 |
| | pPR3-N-MJ-**NOT3 687-844** | pFL-Flag-NOT3 | MS13 |

**REFERENCES**


Jeske, M., Bordi, M., Glatt, S., Muller, S., Rybin, V., Muller, C.W., and Ephrussi, A. (2015). The Crystal Structure of the Drosophila Germline Inducer Oskar Identifies Two Domains with Distinct Vasa Helicase- and RNA-Binding Activities. Cell Rep *12*, 587-598.

Salgania, H.K., Metz, J., and Jeske, M. (2022). ReLo: a simple colocalization assay to identify and characterize physical protein-protein interactions. BioRxiv, doi: 10.1101/2022.1103.1104.482790

Temme, C., Zhang, L.B., Kremmer, E., Ihling, C., Chartier, A., Sinz, A., Simonelig, M., and Wahle, E. (2010). Subunits of the Drosophila CCR4-NOT complex and their roles in mRNA deadenylation. RNA *16*, 1356-1370.
